# The molecular evolution of spermatogenesis across mammals

**DOI:** 10.1101/2021.11.08.467712

**Authors:** Florent Murat, Noe Mbengue, Sofia Boeg Winge, Timo Trefzer, Evgeny Leushkin, Mari Sepp, Margarida Cardoso-Moreira, Julia Schmidt, Celine Schneider, Katharina Mößinger, Thoomke Brüning, Francesco Lamanna, Meritxell Riera Belles, Christian Conrad, Ivanela Kondova, Ronald Bontrop, Rüdiger Behr, Philipp Khaitovich, Svante Pääbo, Tomas Marques-Bonet, Frank Grützner, Kristian Almstrup, Mikkel Heide Schierup, Henrik Kaessmann

## Abstract

The testis is a key male reproductive organ that produces gametes through the process of spermatogenesis. Testis morphologies and spermatogenesis evolve rapidly in mammals, presumably due to the evolutionary pressure on males to be reproductively successful^1,2^. The rapid evolution of the testis was shown to be reflected at the molecular level based on bulk-tissue work^3-8^, but the molecular evolution of individual spermatogenic cell types across mammalian lineages remains largely uncharacterized. Here we report evolutionary analyses of single-nucleus transcriptome data for testes from eleven species that cover the three major mammalian lineages (eutherians, marsupials, egg-laying monotremes) and birds (the evolutionary outgroup), and include seven key primates. Our analyses reveal that the rapid evolution of the testis is driven by accelerated fixation rates of gene expression changes, amino acid altering substitutions, and newly emerged genes in late spermatogenic stages – likely facilitated by reduced pleiotropic constraints, haploid selection, and a transcriptionally permissive chromatin environment. We identify temporal expression changes of individual genes across species, which may have contributed to the emergence of species-specific phenotypes, but also conserved expression programs underlying ancestral spermatogenic processes. Sex chromosome analyses show that genes predominantly expressed in spermatogonia (i.e., germ cells fueling spermatogenesis) and Sertoli cells (i.e., somatic supporting cells) independently accumulated on X chromosomes across mammals during evolution, presumably due to male-beneficial selective forces. Further work uncovered that the process of meiotic sex chromosome inactivation (MSCI) also occurs in monotremes and hence is common to the different mammalian sex chromosome systems, contrary to previous inferences^9^. Thus, the general mechanism of meiotic silencing of unsynapsed chromatin (MSUC), which underlies MSCI, represents an ancestral mammalian feature. Together, our study illuminates the cellular and molecular evolution of mammalian spermatogenesis and associated selective forces, and provides a resource for investigating the biology of the testis across mammals.

---

The rapid evolution of the testis across mammals is likely mainly explained by positive selection associated with the force of sperm competition, which reflects the evolutionary pressure on males to achieve reproductive success^1,2^ (i.e., to fertilize female eggs). Consequently, testis sizes and associated sperm production rates, sperm morphologies, and various other cellular traits substantially vary across mammals and even between closely related species such as the great apes, due to major mating system differences, especially regarding the extent of female promiscuity^1,2^. The rapid phenotypic evolution of the testis is reflected at the molecular level. Previous large-scale gene expression comparisons for various organs across mammals revealed that rates of evolutionary expression change are highest in the testis, probably due to frequent adaptive changes but potentially also widespread relaxation of purifying selection^3-8^. Consistently, genes with testis-specific expression tend to be enriched with genes whose coding sequences have been shaped by positive selection^10,11^. In addition to these frequent genetic alterations of preexisting testis-expressed genes, new genes that emerge during evolution tend to be predominantly expressed in the testis and thus likely also contribute to its rapid phenotypic evolution^5,12^.

The testis displays several other unique molecular features as well. First, chromatin in spermatogenic cells is massively remodeled during spermatogenesis, a process that culminates in the tight packaging of DNA around protamines in the compact sperm head^13^. Notably, this remodeling leads to widespread leaky transcription in the genome^14^, which in turn likely facilitated the initial transcription and – hence – the frequent emergence of new testis-expressed genes and alternative exons during evolution^5,12,14,15^. Second, the differentiation of sex chromosomes from ancestral autosomes triggered the emergence of male meiotic inactivation of sex chromosomes (MSCI) in eutherians and marsupials^16^ (therians), which led to the establishment of backup gene copies that substitute for parental genes on the X during meiosis^5,17^. In spite of MSCI, the X chromosome has become enriched with testis-expressed genes during evolution^5,18-23^, presumably due to sexually antagonistic selective forces favoring the fixation of male-beneficial mutations on this chromosome^24^. Finally, translational regulation of transcriptomes is particularly intricate and widespread across spermatogenesis^8^.

Previous large-scale transcriptomic investigations of testis evolution were largely limited to bulk-organ samples^3-8,22^. Recent high-throughput single-cell (sc) or single-nucleus (sn) RNA sequencing (RNA-seq) technologies^25^ enable detailed investigations of the cellular and molecular evolution of the testis, as exemplified by two scRNA-seq comparisons between human, macaque, and mouse^26,27^, but a comprehensive investigation of the evolution of spermatogenesis across all major mammalian lineages is lacking.

Here we provide an extensive snRNA-seq resource covering testes from ten key representative mammalian species and a bird, enabling detailed comparisons of spermatogenic cells and underlying gene expression programs within and across mammals (https://apps.kaessmannlab.org/SpermEvol/). Our evolutionary analyses of these data unveiled the ancestral as well as species- and lineage-specific cellular and molecular characteristics of mammalian spermatogenesis.

## Spermatogenesis across eleven species

We generated snRNA-seq data for the testis from ten species that cover the three major mammalian lineages and include key primate species, representing all simian (anthropoid) lineages (Fig. 1a): eutherian mammals (representatives for 5 of the 6 extant ape lineages, including humans; rhesus macaque, an Old World monkey; common marmoset, a New World monkey; mouse), marsupials (grey short-tailed opossum), and egg-laying monotremes (platypus). Corresponding data were generated for a bird (red junglefowl, the progenitor of domestic chicken; hereafter referred to as ‘chicken’), to be used as an evolutionary outgroup. The dataset overall consists of 34 libraries, with one to three biological replicates per species. We refined and extended existing Ensembl^28^ genome annotations across all species based on RNA-seq data generated for bulk-testis samples (Supplementary Tables 1, 2; Methods), to ensure optimal read-mapping and prevent biases in downstream cross-species analyses. After quality controls and filtering steps (Methods), we obtained transcriptomes for a total of 97,521 high-quality nuclei for the eleven species (Supplementary Tables 2, 3).

**Fig. 1.**
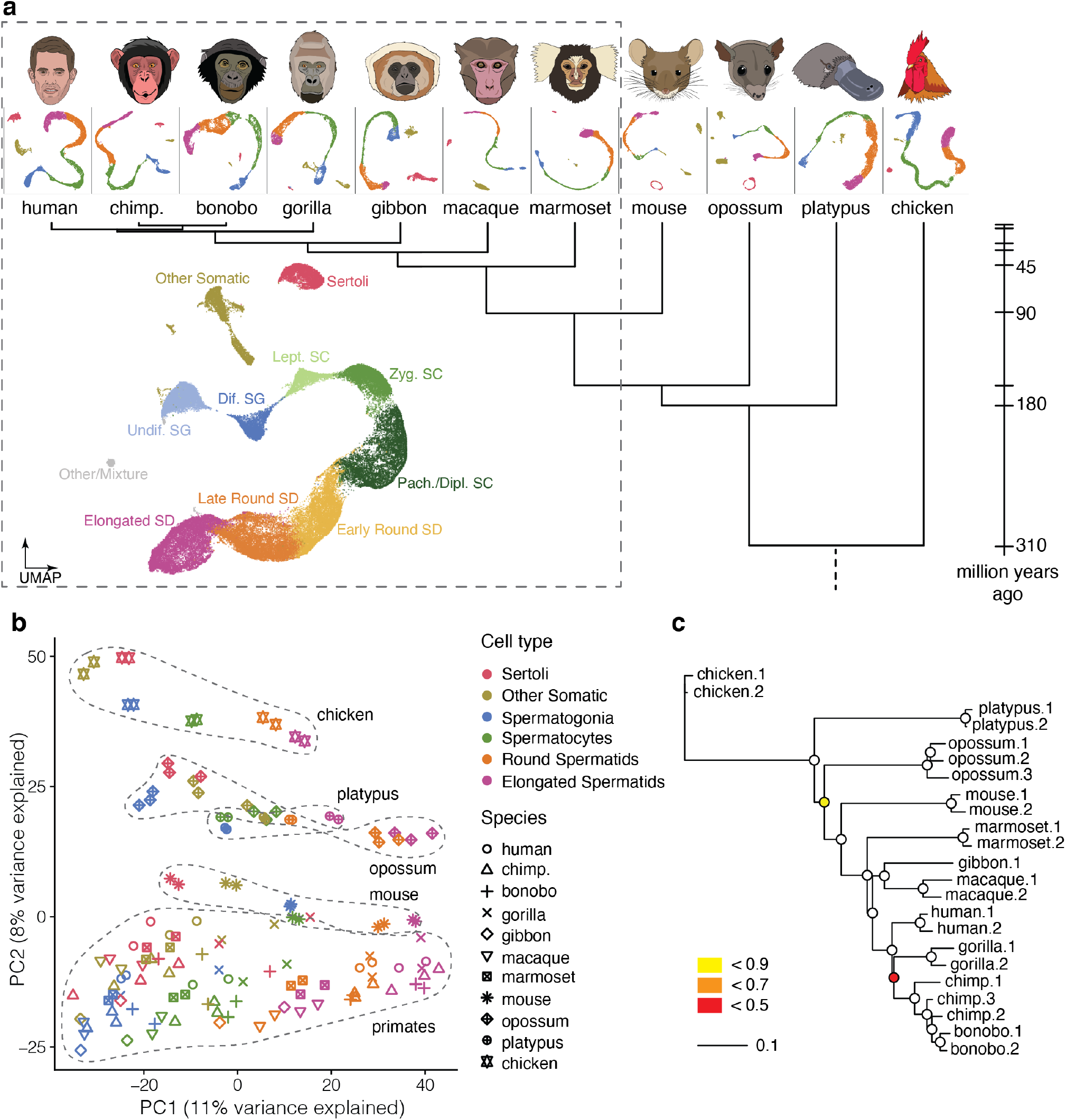
Single nucleus RNA profiling across 10 mammals and a bird. **a**, Species sampled and uniform manifold approximation and projection (UMAP) of snRNA-seq datasets. UMAP of the integrated primate dataset (dashed box), showing undifferentiated and differentiated spermatogonia (Undif. SG and Dif. SG, respectively), leptotene, zygotene, pachytene, and diplotene spermatocytes (Lept. SC, Zyg. SC, Pach. SC, and Dipl. SC, respectively), spermatids (SD), and somatic cell types. **b**, Principal component (PC) analysis of cell type pseudo-bulks based on 4,498 1:1 orthologues. Species/lineages are encircled by a dashed line. Each symbol represents an individual. **c**, Gene expression phylogeny based on pseudo-bulk transcriptomes for whole testes. Bootstrap values (4,498 1:1 orthologous amniote genes were randomly sampled with replacement 1,000 times) are indicated by circles, ≥ 0.9 (white fill).

Using a clustering approach and marker gene based annotation procedure (Methods), we identified the main germ cell types^29^ along the continuous cellular proliferation and differentiation path of spermatogenesis across all species: spermatogonia (SG), the mitotic cells fueling spermatogenesis and including spermatogonial stem cells; spermatocytes (SC), where meiosis takes place; and the haploid round spermatids (rSD) and elongated spermatids (eSD), which together reflect the process of spermiogenesis (Fig. 1a; Extended Data Figs. 1a-f; Supplementary Table 3; Methods). We also identified separate clusters corresponding to somatic testicular cells, in particular Sertoli cells (except for platypus, in which Sertoli cells could not be unambiguously distinguished), the main spermatogenesis support cells, but also other cell types, such as Leydig cells, peritubular cells, endothelial cells and macrophages (Extended Data Figs. 1a-e). The close evolutionary relationship of the seven primates in our study, which diverged between ∼2-43 million years ago, enabled the direct integration of datasets across these species and thus the identification of sub-cell types that correspond to intermediate events during spermatogenesis (Fig. 1a). For all species, we traced the highly dynamic gene expression programs underlying spermatogenic differentiation and key molecular events, thus also identifying a host of novel marker genes (Extended Data Fig. 1g; Supplementary Table 4).

To obtain an initial overview of cell type relationships across species, we performed a principal-component analysis based on pseudo-bulk cell type transcriptomes (Fig. 1b). The first principal component (PC1), explaining most variance in gene expression, orders the spermatogenic cell types according to the progression of spermatogenesis for all species (Fig. 1b, from left to right). This observation suggests that our data capture ancestral aspects of spermatogenic gene expression programs that are shared across mammals/amniotes, despite the rapid evolution of the testis^3-8^ and the long divergence times of 310 million years (Fig. 1a). PC2 separates the data by species or lineages, reflecting diverged aspects of spermatogenesis, while PC1 and PC2 together separate somatic and spermatogenic cell types for each species. The consistent and close clustering of samples from biological replicates is a further indicator of the high data quality.

## Rates of evolution along spermatogenesis

To investigate the rates of gene expression evolution across testicular cell types, we reconstructed gene expression trees (Methods). A tree based on pseudo-bulk transcriptomes for the whole testis (Fig. 1c) recapitulates well the known mammalian phylogeny (Fig. 1a; except for the gibbon-macaque grouping), akin to previous trees based on bulk-tissue RNA-seq data across mammalian organs, which originally revealed the rapid expression evolution of the mammalian testis^3,4^. This observation is consistent with the view that regulatory changes steadily accumulated over evolutionary time^3^, with present-day RNA abundances reflecting the evolution of mammalian lineages and species.

To trace the cellular source of the rapid evolution of the testis, we next built expression trees for the different testicular cell types, which also recapitulate well the known species relationships (Extended Data Fig. 2). Notably, the total branch lengths of the trees, which reflect the amount of evolutionary expression change, substantially vary between certain cell types (Fig. 2a). While the rate of expression evolution is similar in Sertoli cells and diploid spermatogenic cells (and lower than that in other somatic cell types), it is substantially higher in the post-meiotic haploid cell types (i.e., in rSD and eSD), consistent with a recent inference based on data for three eutherians^27^. The higher resolution afforded by a primate-specific analysis provides further details (Fig. 2a); starting in late meiosis (pachytene SC), evolutionary rates progressively increase until the end of spermiogenesis (late eSD). Thus, late spermatogenic stages drive the previously observed rapid evolution of the testis^3-8^.

**Fig. 2.**
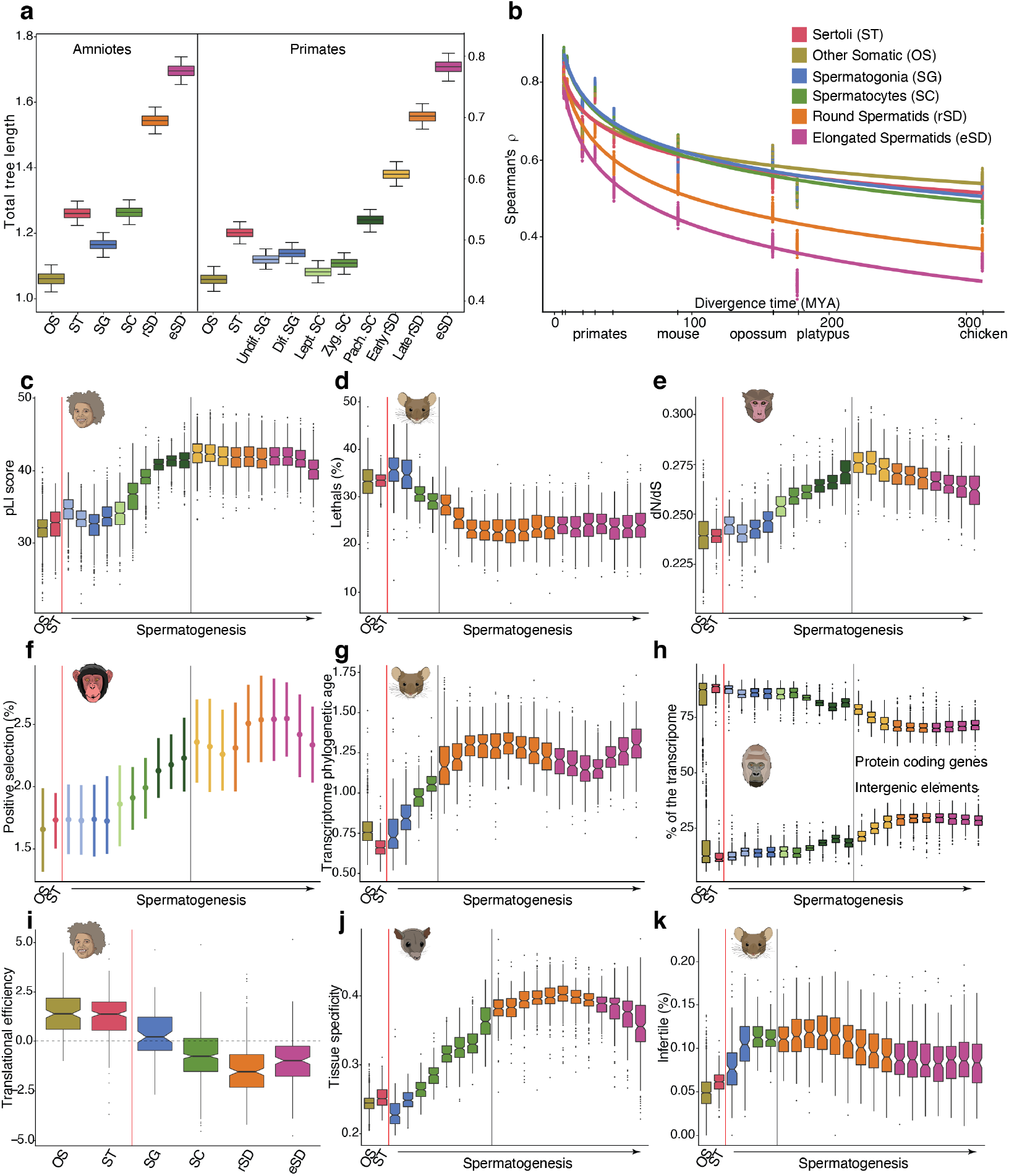
Gene expression divergence and evolutionary forces. **a**, Comparisons of total branch lengths of expression trees among the six main testicular cell types for amniotes and primates. Errors, 95% confidence intervals based on bootstrapping analyses (1,000 replicates). **b**, Spearman’s correlations between humans and other species. Dots correspond to values obtained in 100 bootstrap replicates. The lines correspond to linear regression trends (after log transformation of the time axis) and were added for visualization purposes. Regression R^2^ values range from 0.86 to 0.97. **c**, Tolerance to functional variants in human (the pLI score^30^ reflects the tolerance of a gene to a loss-of-function mutation; lower values mean less tolerance). **d**, Percentage of expressed genes that lead to a lethal phenotype when knocked out in mouse (out of 4,742 knockouts). **e**, Average normalized ratio of nonsynonymous (dN) over synonymous (dS) nucleotide substitutions across primates (data shown for macaque; other species in Extended Data Fig. 3b). **f**, Percentage of positively selected genes (out of 11,170 genes tested for positive selection; data shown for chimp, other species in Extended Data Fig. 3c). Whiskers depict the interquartile ranges. **g**, Phylogenetic age of cellular transcriptomes (data shown for mouse, other species in Extended Data Fig. 3d). Higher values reflect increased lineage-specific gene contributions (that is, younger transcriptomes). **h**, Percentage of UMIs from protein coding genes (top) or intergenic elements (bottom) (data shown for gorilla, other species in Extended Data Fig. 3e). **i**, Translational efficiency (TE; normalized log2-transformed values; data shown for human, other species in Extended Data Fig. 3f). **j**, Tissue specificity (data shown for opossum, other species in Extended Data Fig. 3g). **k**, Percentage of mouse genes associated with infertility (out of 3,252 knockouts). In **c, j** and, **e, g**, reported values correspond to mean and to the median values across genes expressed in each cell, respectively. In **c-e, g-k**, box plots depict the interquartile ranges, with whiskers at 1.5 times the interquartile range. In **c-e, g, h, j, k**, germ cells are binned into 20 equally sized groups progressing through spermatogenesis using pseudotime trajectories (from left to right). The gray vertical line denotes the transition from diploid spermatocytes to haploid spermatids. In **c**-**k**, cells are grouped by cell type with somatic cells left of red vertical line.

Pairwise species comparisons confirm the rapid expression evolution of post-meiotic cell types across amniotes and that gene expression divergence overall increases with evolutionary time (Fig. 2b), in accord with the expression phylogeny results (Extended Data Fig. 2). It is noteworthy, however, that expression divergence levels are approximately as similar between human and chicken as they are between human and platypus, although the bird lineage diverged 110 million years before the separation of monotremes and therian mammals (i.e., eutherians and marsupials). This observation had been made for whole organs^3^ and supports at the cellular level the notion that the conservation of core spermatogenic functions restricts transcriptome divergence.

## Evolutionary forces

We next sought to trace the evolutionary forces underlying the rapid evolution of late spermatogenesis. Two non-mutually exclusive patterns of natural selection may account for this observation. First, later stages of spermatogenesis might evolve under weaker purifying selection (i.e., reduced functional constraints) and hence be less refractory to change. Second, the greater divergence in later stages might result from stronger positive selection, increasing the rate of fixation of adaptive changes. To investigate patterns of functional constraint during spermatogenesis, we assessed the tolerance to functional mutations of genes used in different spermatogenic stages using dedicated human data^30,31^, which revealed a progressive increase of mutational tolerance starting during meiosis and culminating in early spermiogenesis (Fig. 2c). Consistently, based on a set of neutrally ascertained mouse knockouts^32^, we found that the percentage of expressed genes associated with lethality decreases during spermatogenesis (Fig. 2d). Also, in agreement with a progressive reduction in functional constraints towards later spermatogenic stages, we find that the normalized rate of amino acid altering substitutions in coding sequences across primates is higher in late spermatogenesis (Fig. 2e; Extended Data Fig. 3b), although this increase might also reflect a higher proportion of genes under positive selection. Indeed, an examination of the temporal expression pattern of genes whose encoded protein sequences have been shaped by positive selection revealed a significant increase in percentages of positively selected genes used during spermatogenesis, with a peak in rSD (Fig. 2f; Extended Data Fig. 3c).

Given that new genes also contribute to adaptive evolutionary innovations, we investigated the temporal distribution of genes that recently emerged through gene duplication to gene expression programs in germ cells, using an index that combines the phylogenetic age of genes with their expression^4^ (Methods). This analysis revealed that transcriptomes become younger during spermatogenesis (Fig. 2g; Extended Data Fig. 3d), indicating that new duplicate genes have increasingly more prominent roles in later stages, in particular in rSD, consistent with previous observations for human/mouse-specific genes^26,27^. It is noteworthy in this context that previous work based on bulk cell type analyses in mouse^14^ uncovered an overall transcriptionally permissive chromatin environment during spermatogenesis, in particular in rSD, which was suggested to have facilitated the emergence of new genes during evolution^5,12,14^. Consistently, we detect, in all species, significantly increased contributions of intergenic transcripts to the transcriptomes after meiosis and a concomitant decrease in the contributions of protein-coding genes (Fig. 2h; Extended Data Fig. 3e). Notably, based on analyses integrating previous translatome data^8^, we detected a decline of translational efficiencies of transcripts during spermatogenesis (reaching a minimum in rSD) in all studied species (Fig. 2i; Extended Data Fig. 3f). This decline is consistent with previous observations from mouse bulk data for a restricted number of cell types^8^ and likely mitigates the functional consequences of the concurrent increase in transcriptional promiscuity of the genome.

We next explored the reasons underlying the dynamic changes of selective forces and patterns of innovation during spermatogenesis. The breadth of expression across tissues and developmental processes (here referred to as pleiotropy) was shown to represent a key determinant of the types of mutations that are permissible under selection^33,34^. We, therefore, assessed patterns of pleiotropy across spermatogenesis through analyses that incorporated spatiotemporal transcriptome data for mammalian organs^4^, which revealed that genes used later in spermatogenesis, in particular those in rSDs, have substantially more specific spatial and temporal expression profiles than genes employed earlier in spermatogenesis and in somatic cells (Fig. 2j; Extended Data Fig. 3g, h). Given that a decrease in pleiotropy can explain both a decrease in functional constraints and an increase in adaptation^4,33,34^, we suggest that it is likely to be a major contributor to the accelerated molecular evolution observed across species in late spermatogenesis. In addition, the specific type of selection acting on haploid cells^35,36^ (haploid selection), where expressed alleles are directly exposed to selection, may have contributed to the exceptionally rapid evolution of the haploid rSD.

While the very tissue- and time-specific late spermatogenic genes, in general, are not essential for viability (Fig. 2d, j; Extended Data Fig. 3g, h; see above), we hypothesized that the specific aforementioned evolutionary forces imply that many of these genes evolved crucial roles in spermatogenesis, such as spermatid-specific protamine-encoding genes^37^. Indeed, we find that the proportion of genes associated with infertility^32^ is relatively high in SC and SDs (especially rSDs) – higher than in SG and somatic testicular cells (Fig. 2k).

## Evolution of gene expression trajectories

We next sought to trace the individual genes underlying conserved (ancestral) and diverged aspects of germ cells by comparing expression trajectories along spermatogenesis of 1:1 orthologous genes across species using a previously established phylogenetic procedure^4^ (Extended Data Fig. 4; Methods). For the primates, this analysis revealed between ∼1,700 and ∼2,900 genes with conserved expression trajectories (e.g., *TEX11*) across different lineages or species (Fig. 3a; Supplementary Table 5). For example, the temporal expression patterns of 1,687 genes are conserved across the seven primates and hence likely reflect the core of the ancestral gene expression program of the simian testis.

**Fig. 3.**
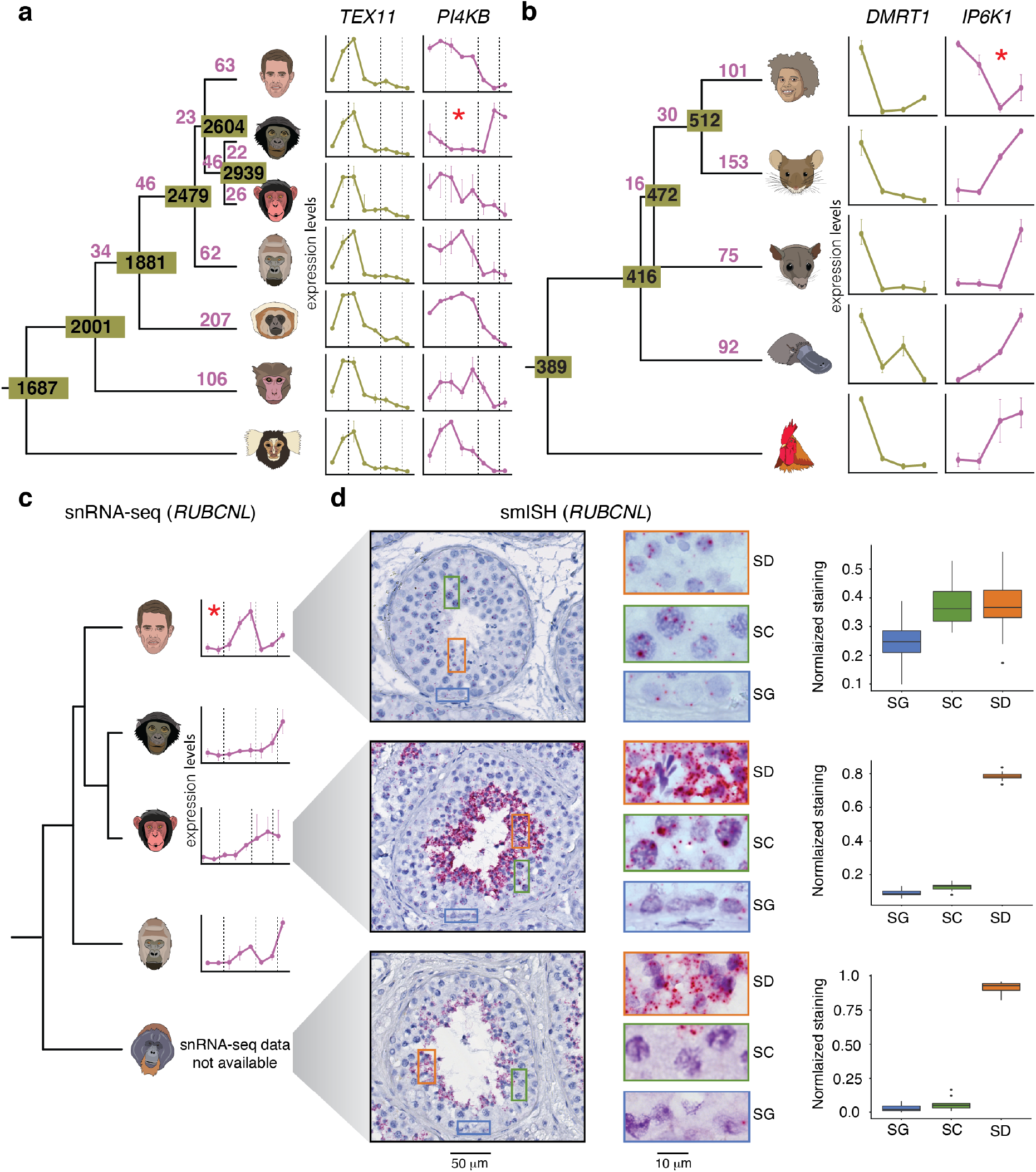
Evolution of gene expression trajectories along spermatogenesis. **a** and **b**, Numbers depict the amount of trajectory changes (in purple) and conserved trajectories (in olive). In **a, b**, and **c**, red asterisks indicate the branch for which a trajectory change has been called. Trajectories show expression levels along spermatogenesis (from left to right). In **a** and **c**, the dotted vertical lines separate spermatogonia (SG), spermatocytes (SC), round (rSD) and elongated spermatids (eSD) from left to right. **a**, Primate trajectories (human, bonobo, chimp., gorilla, gibbon, macaque, and marmoset; from top to bottom) based on 4,459 1:1 orthologues. *TEX11* and *PI4KB* are examples of conserved and changed trajectories, respectively. **b**, Amniote trajectories (human, mouse, opossum, platypus, and chicken; from top to bottom) based on 2,927 1:1 orthologues. *DMRT1* and *IP6K1* are examples of conserved and changed trajectories, respectively. **c**, *RUBCNL* expression trajectory from snRNA-seq along spermatogenesis for human, bonobo, chimpanzee, and gorilla (from top to bottom). Available orangutan samples did not allow for the generation of high-quality snRNA-seq data. **d**, Detection of *RUBCNL* (each red dot corresponds to a single transcript) expression in human, chimpanzee, and orangutan testis by smISH using RNAScope (from top bottom). Left: Seminiferous tubule cross sections counterstained with hematoxylin, and closeups on areas containing SG, SC, and SD. Right: Quantification of *RUBCNL* expression levels in 10 tubules per section (n = 3 for human, n = 1 for chimp. and orangutan). Box plots represent the distribution of staining, with whiskers at 1.5 times the interquartile range.

By contrast, using the marmoset as an outgroup, we detected 635 trajectory changes during the evolution of apes and Old World monkeys (catarrhines) (Fig. 3a; Supplementary Table 5). For example, 63 and 94 trajectory changes occurred on the human and chimpanzee/bonobo lineages, respectively, since their divergence ∼7 million years ago (e.g., *PI4KB*). We sought to spatially validate three of these changes using single-molecule RNA *in situ* hybridization (smISH) for three key great ape species – human, chimpanzee, and orangutan (the outgroup) – for which suitable samples could be sourced (Fig. 3c; Extended Data Fig. 5, 6; Supplementary Table 6; Method). In accord with the trajectory analyses, our smISH experiments confirm that the expression of the gene *RUBCNL*, encoding a regulator of autophagy^38^, changed on the human lineage, with an overall relative reduction of expression levels in SD compared to chimpanzee and orangutan (Fig. 3c, d). Thus, the functional role of *RUBCNL* in autophagy during spermatogenesis, where this process is involved in a broad range of cellular events^39^, is potentially different in humans compared to other catarrhines. We also confirmed the inversion of the expression trajectory of the myosin-encoding gene *MYO3B* in SG and rSD on the chimpanzee/bonobo lineage (Extended Data 5a), as well as the expression increase of *ADAMTS17* – encoding a family-member of proteases with key spermatogenic functions^40^ – in rSD relative to SG in humans (Extended Data Fig. 5b). We note that observed quantitative and partly qualitative expression pattern differences between the two complementary data types are expected because of technical differences. For example, the trajectory analyses reflect transcript abundances in nuclei, whereas smISH allows quantification in whole cells. On the other hand, substantial nucleus condensation limits smISH-based transcript quantification in eSD.

We also assessed the conservation of expression trajectories across the three major mammalian lineages and amniotes as a whole (Fig. 3b; Supplementary Table 7). For example, the 416 genes with conserved expression across mammals likely trace back to ancestral gene expression programs of the ancestral mammalian testis, while 389 genes may represent the core ancestral spermatogenic program of amniotes. A notable example is *DMRT1*, which is highly expressed specifically in SG across amniotes and is required for mouse spermatogonial stem cell maintenance and replenishment^41^.

In agreement with the notion that these highly conserved sets of genes play key roles in mammalian/amniote spermatogenesis, our analyses of fertility phenotypes^32^ unveiled that genes involved in fertility are overall significantly more conserved in their expression trajectories than genes not associated with fertility (Extended Data Fig. 7b). Thus, genes with conserved trajectories for which spermatogenesis functions remain uncharacterized represent a promising resource of candidates for the exploration of fertility phenotypes (Supplementary Tables 5, 7). Notably, however, also among genes with lineage-specific trajectory changes, we identified genes for which key fertility function have been described in mouse or human. For example, *IP6K1*, which elicits infertility when knocked out in the mouse, shows strongly increasing expression towards the end of spermiogenesis in all amniotes except human and the other primates, where expression is high in SG and then overall declines (Fig. 3b; Extended Data Fig. 7c). This suggests that the primary function of *IP6K1* may have shifted from late to early spermatogenesis (SG) during primate evolution.

## Sex chromosomes

Sex chromosomes emerged twice in parallel in mammals from different sets of ancestral autosomes. The therian XY chromosome system originated in the common therian ancestor just prior to the split of eutherians and marsupials (Fig. 4a) and hence has evolved largely independently in these two lineages. Around the same time, the original pair of the monotreme sex chromosome system arose from different autosomes and subsequently expanded during evolution to the five XY pairs seen in monotremes today^42^. These sex chromosome formation events entailed substantial remodelling of gene contents and expression patterns due to structural changes and sex-related selective forces^5^. We used our data to systematically assess testicular expression patterns of sex chromosomal genes and their evolution across mammals.

**Fig. 4.**
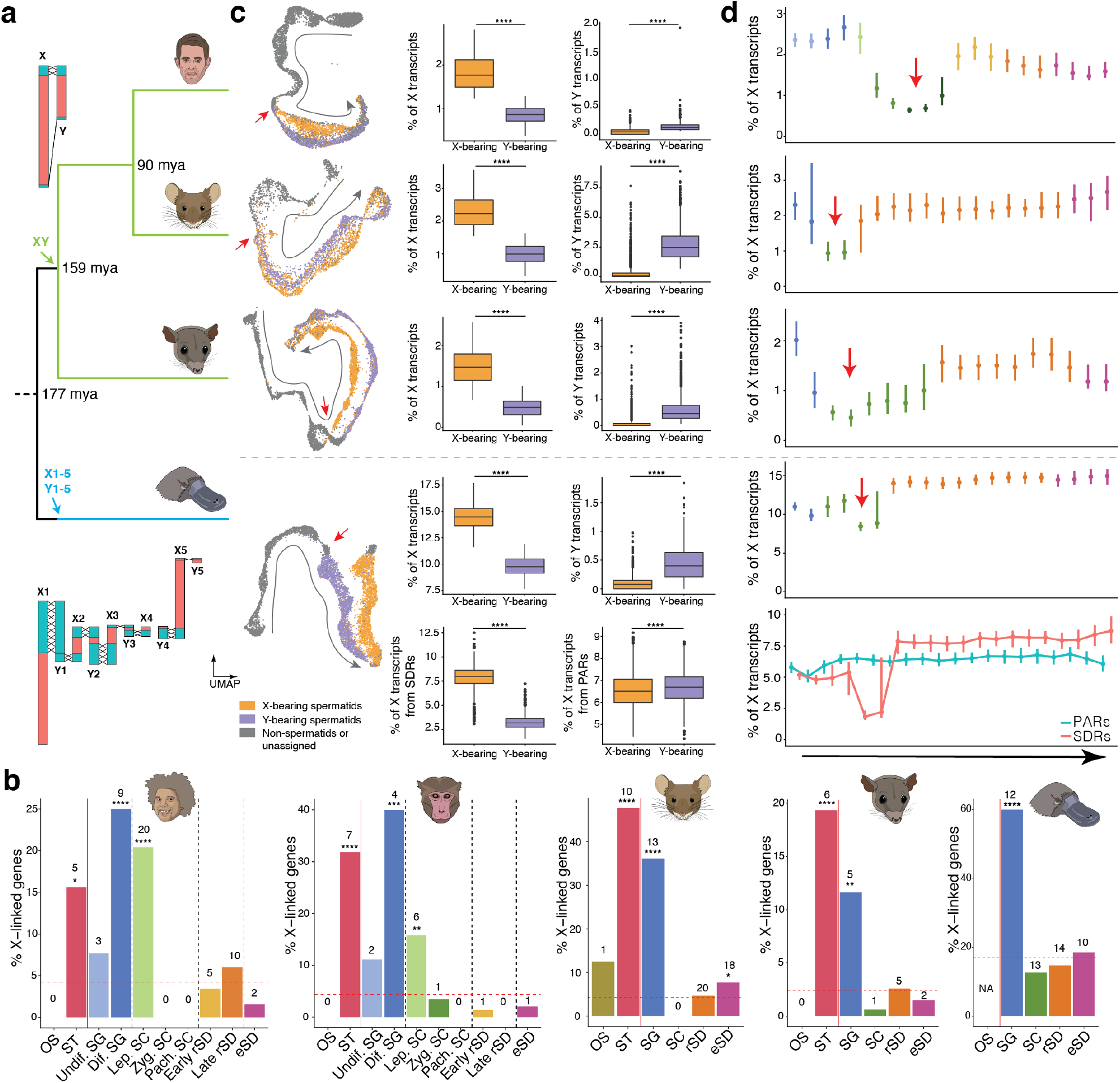
Mammalian sex chromosome evolution. **a**, Ages of main speciation events obtained from TimeTree (www.timetree.org), for human, mouse, opossum, and platypus (from top to bottom), are indicated (millions of years ago, mya). Arrows show when sex systems originated in mammals^42^. Illustration of therian (top) and monotreme (bottom) sex chromosomes, using human^45^ and platypus^46^ as representatives, respectively. Pseudo-autosomal regions (PARs) in turquoise and sexually differentiated regions (SDRs) in red. Crosses indicate recombining regions. **b**, Percentages of X-linked genes among testis-specific genes with predominant expression in each cell type, for human, macaque, mouse, opossum, and platypus (from left to right). The red horizontal dashed line represents the expected percentages of X-linked genes, if testis-specific genes with predominant expression in the different cell types were randomly distributed across the genome. Exact binomial tests were performed for statistical comparisons (\**P* < 0.05, \*\**P* < 0.01, \*\*\**P* < 0.001, \*\*\*\**P* < 0.0001). The number of testis-specific X-linked genes enriched in each cell type is indicated above each bar. **c**, UMAP representation of germ cells (left). The grey arrows indicate the progression of spermatogenesis. The red arrows indicate the meiotic divisions. Spermatids identified as X and Y-bearing cells are colored in orange and purple, respectively. Box plots show the percentages of X and Y transcripts in X and Y-bearing spermatids. For platypus, X transcripts are dissected according to their location on the X chromosomes (PARs and SDRs annotated by ref.^46^). In the platypus genome assembly used^46^, X and Y PARs are assigned to the X chromosome. Thus, reported X transcripts can come from X SDRs, X PARs or Y PARs, while reported Y transcripts only from Y SDRs. Two-sided Wilcoxon rank-sum tests were performed for statistical comparisons (\**P* < 0.05, \*\**P* < 0.01, \*\*\**P* < 0.001, \*\*\*\**P* < 0.0001). **d**, Median percentages of X transcripts in germ cells grouped into 20 equally sized bins across spermatogenesis (from left to right). For spermatids, only X-bearing cells are considered. For platypus, X transcripts are dissected according to their location on the X chromosomes (PARs and SDRs). Red arrows indicate MSCI. Whiskers depict the interquartile ranges.

We first contrasted cell type specificities of X-linked and autosomal genes, which revealed a striking excess (12-60%) of genes with predominant expression in SG uniquely located on the X chromosome across all eutherians, which is consistent with a previous mouse study^20^, and includes conserved genes with key spermatogenic functions such as *TEX11* (Fig. 3a; Fig. 4b; Extended Data Fig. 8a; Supplementary Table 8). The high-resolution primate data revealed that the X chromosome is also enriched for genes with expression in leptotene SC (Fig. 4b), presumably reflecting transcript carry-over from SG, given the global transcriptional silencing of the genome during the leptotene stage of spermatogenesis^43,44^. Here we show that this genome-wide transcriptional silencing represents a shared and thus likely ancestral feature of amniotes (Extended Data Fig. 9; Supplementary Table 3). In agreement with the notion of a transcript carry-over from SG, most of the X-linked genes expressed in leptotene SC are also expressed in differentiated SG (Extended Data Fig. 10). Notably, we detected enrichments of genes with SG-specific expression also on the opossum and platypus X chromosomes, thus revealing this to be a shared pattern across mammals (Fig. 4b; Extended Data Fig. 8b).

Additionally, our analyses uncovered an enrichment of genes with predominant expression in Sertoli cells across mammalian X chromosomes (except for platypus, where Sertoli cells could not be unambiguously distinguished; see above) (Fig. 4b). Altogether, our observations suggest that sex-related forces favored the independent accumulation of SG- and Sertoli-specific genes on the X during the evolution of the different mammalian sex chromosome systems. In support of this view, autosomes in outgroup species corresponding to the different mammalian X chromosomes, (e.g., platypus chromosome 6, which is homologous to the therian X), do not show any excess of SG- /Sertoli-expressed genes (Extended Data Fig. 8a, c); that is, ancestral autosomes that gave rise to present-day sex chromosomes were likely not yet enriched for such genes. Overall, we hypothesize that the accumulation of SG/Sertoli-specific genes was facilitated by the specific selective environment on the X chromosome upon sex chromosome differentiation from ancestral autosomes, in which male-beneficial mutations are always visible to positive selection because of the single-copy (that is, hemizygous) status of the X chromosome in this sex^5,20,24^.

We next sought to investigate the expression of sex chromosomal genes across spermatogenesis – separately for X- and Y-linked genes. While such an analysis is likely not possible using single-whole-cell transcriptomic data, given that X and Y spermatids remain connected by cytoplasmic bridges and hence are thought to contain largely similar cytoplasmic transcript pools^47,48^, our single-nucleus data should afford the separation of X/Y spermatids. Indeed, based on differential X/Y transcript contents across spermatids (Methods), we were able to separate spermatids into distinct X and Y lineages across mammals (Fig. 4c; Supplementary Table 3).

Based on these data, we observed a substantial dip in X transcript abundances around the pachytene stage of meiosis across therians (Fig. 4d), in accord with previous work^16,49^, reflecting the process of MSCI – a sex chromosome-specific instance of the general epigenetic phenomenon of meiotic silencing of unsynapsed chromatin^16,49^ (MSUC). An analysis of Y transcripts provides consistent results, albeit at much lower resolution, due to the very small number of Y-linked genes (Extended Data Fig. 11a). Previous work did not reveal any evidence for MSCI in monotremes^9^, leading to the suggestion that this process originated in the therian ancestor; that is, after the separation of the monotreme lineage from that of therians^9^. We revisited this question using our unique platypus data and a new high-quality platypus genome assembly and annotation^46^ that includes, for the first time, a detailed definition of sexually differentiated regions (SDRs) versus pseudoautosomal regions (PARs), which turn out to be large in monotreme sex chromosomes (Fig. 4a). We hypothesized that expression signals from the large PARs, which are expected to synapse and hence not to be affected by MSUC, might have prevented the detection of MSCI in previous transcriptomic studies.

Indeed, while the joint analysis of all platypus X-linked genes only reveals a small potential expression dip around the pachytene stage (Fig. 4d, upper platypus graph), a focused analysis of SDR genes reveals a strong reduction of X transcript levels. By contrast, PAR genes show stable expression levels across spermatogenesis (Fig. 4d, lower graph). Moreover, the difference in transcript abundances between SDRs and PARs due to MSCI is also clearly visible for all of the five platypus X chromosomes when assessing average expression levels along the chromosomes (Extended Data Fig. 11b). Notably, the presence of MSCI at the SDRs in monotremes is consistent with the previously observed partial association of platypus sex chromosomes with perinucleolar repressive histone modifications at the pachytene stage^9^. Altogether, our data reveal that MSCI is common to mammalian sex chromosome systems, which implies that the general mechanism of MSUC represents an ancestral mammalian feature.

## Discussion

In this study, we used extensive snRNA-seq testis data from ten representative mammals and a bird, as well as complementary data (e.g., smISH, translatome data), to scrutinize the evolution of mammalian spermatogenesis. Our analyses uncovered that the previously observed rapid evolution of the testis at both the phenotypic^1,2^ and molecular level^3-8^ is driven by an accelerated rate of molecular innovation in late spermatogenesis, in particular in rSD cells. Specifically, our findings suggest a scenario where the accelerated fixation of regulatory changes, amino acid altering substitutions, and new genes during evolution in late spermatogenesis – driven by sperm competition (positive selection) – was facilitated by reduced pleiotropic constraints, the haploidy of spermatids (i.e., haploid selection), and a transcriptionally permissive chromatin environment. Consistently, many genes expressed at the late stages of spermatogenesis are associated with infertility while only few are essential, suggesting that they overall evolved important spermatogenic functions under the aforementioned evolutionary pressures. In agreement with the notion that late spermatogenic stages explain the rapid evolution of the testis observed at the bulk-tissue level, early spermatogenic cell types and somatic testis cells show patterns of constraint and innovation that are similar to those of cell types in the brain (Extended Data Fig. 3a, b, d), an organ that evolves overall slowly at the molecular level^3-5^. We note, however, that differences in cell type abundances across mammals, which are pronounced for the testis^1^ (Extended Data Fig. 1f), presumably also contributed to the previously observed rapid gene expression divergence of this organ^8^.

Our cross-species comparisons of individual genes revealed temporal expression differences across species, including human-specific changes, which probably contributed to species-specific spermatogenesis phenotypes, as well as conserved expression programs underlying spermatogenic processes ancestral to individual mammalian lineages and mammals as a whole.

Our analyses also illuminated the role of sex chromosomal genes in spermatogenesis. We found that genes predominantly expressed in spermatogonia (i.e., germ cells fueling spermatogenesis) and Sertoli cells (i.e., somatic support cells) independently accumulated on X chromosomes across mammals and their two distinct sex chromosome systems during evolution. This suggests that mammalian X chromosomes have been shaped by strong male-related (beneficial) selective forces^24^, leading to the emergence of many X-linked genes with functional roles in testis cell types where active transcription is possible. Indeed, in addition to the SG/Sertoli-cell enrichment found here, previous work showed that these selective pressures led to the repeated duplication of genes on the X that facilitated expression after meiosis^18^. Further expression analyses across spermatogenesis, which benefited from the separation of X- and Y-bearing spermatids and a new high-quality platypus genome assembly^46^, unveiled that MSCI is a general feature of mammalian sex chromosome systems, implying that MSUC was present in the common mammalian ancestor. Previous work did not find evidence for MSCI in birds^50^, which raises the question of whether MSUC arose in the common mammalian ancestor after the split from the avian/reptile lineage ∼300 million years ago, or whether this mechanism was lost on that lineage and arose earlier in evolution. The latter scenario would be consistent with the observation of MSCI in several invertebrate species^51-53^.

Altogether, our study illuminates the cellular and molecular basis of the evolution of spermatogenesis and associated selective forces across all major mammalian lineages and a range of key primate species. Our data and results, together with the accompanying online resource we have developed (https://apps.kaessmannlab.org/SpermEvol/), which allows gene expression profiles of orthologous genes to be explored across the studied species, provide an extensive resource for investigating the biology of the testis and associated fertility disorders across mammals.

## Acknowledgements

We thank all members of the Kaessmann group, Stephan Tirier for discussions, and Nils Trost for the administration of the Kaessmann lab server. Computations were performed on the Kaessmann lab server and the bwForCluster from the Heidelberg University Computational Center (supported by the state of Baden-Württemberg through bwHPC and the German Research Foundation - INST 35/1134-1 FUGG). This research was supported by grants from the European Research Council (615253, OntoTransEvol) and German Research Council (DFG, SFB 873) to H.K., by the CellNetworks Postdoc Fellowship and EMBO Long-Term Fellowship to F.M. (ALTF 591-2017), and by the Australian Research Council (FT160100267) to F.G.

## Author contribution

F.M., N.M. and H.K. conceived and organized the study based on H.K.’s original design. F.M., N.M. and H.K. wrote the manuscript, with input from all authors. F.M. performed all analyses, developed the shiny app, and drew the species icons. K.M., T.B. collected the samples. R.B., P.K, F.G, S.P., I.K., Ro.B., T.M.B., K.A. and M.H.S. provided samples. F.M., N.M., T.T., and M.S. established snRNA-seq methods. N.M. performed the snRNA-seq experiments with support from J.S., C.S., K.M., and T.B.. S.B.W. performed the smISH experiments, and analyzed the results together with K.A.. E.L. performed the testis genomic annotations. M.S., M.C.M, F.L., M.R.B., C.C., K.A., and M.H.S. provided useful feedback and discussions.

## Correspondence and requests for materials

should be addressed to F.M., N.M., or H.K.

## Methods

### Data reporting

No statistical methods were used to predetermine sample size. The experiments were not randomized and investigators were not blinded to allocation during experiments and outcome assessment.

### Biological samples and ethics statement

We generated single-nucleus RNA-seq data for adult testis samples from human (*Homo sapiens*), chimpanzee (*Pan troglodytes*), bonobo (*Pan paniscus*), gorilla (*Gorilla gorilla*), lar gibbon (*Hylobates lar*), rhesus macaque (*Macaca mulatta*), common marmoset (*Callithrix jacchus*), mouse (*Mus musculus*, strain: RjOrl:SWISS; Janvier Labs), grey short-tailed opossum (*Monodelphis domestica*), platypus (*Ornithorhynchus anatinus*), and chicken (red jungle fowl, *Gallus gallus*) (Supplementary Table 2). In addition, we produced bulk RNA-seq data for chimpanzee, gorilla, gibbon, and marmoset from the same individuals. Adult human testis samples used for *in situ* hybridization experiments were obtained from orchiectomy specimens from three individuals with testicular cancer. Tissue adjacent to the tumor that was devoid of cancer cells and germ cell neoplasia *in situ*, and tubules with normal spermatogenesis were used. Other adult primate testis tissue was obtained from a western chimpanzee and a Bornean orangutan (*Pongo pygmaeus*).

Our study complies with all relevant ethical regulations with respect to both human and other species’ samples. Human samples were obtained from scientific tissue banks or dedicated companies; informed consent was obtained by these sources from donors before death or from next of kin. The use of all human samples for the type of work described in this study was approved by an ethics screening panel from the European Research Council (ERC) (associated with H.K.’s ERC consolidator grant 615253, OntoTransEvol) and local ethics committees: from the Cantonal Ethics Commission in Lausanne (authorization 504/12); from the Ethics Commission of the Medical Faculty of Heidelberg University (authorization S-220/2017); and from the regional medical research ethics committee of the capital region of Copenhagen (H-16019637). All primates used in this study suffered sudden deaths for reasons other than their participation in this study and without any relation to the organ sampled. The use of all other mammalian samples for the type of work described in this study was approved by ERC ethics screening panels (ERC starting grant 242597, SexGenTransEvolution; and ERC consolidator grant 615253, OntoTransEvol).

### Nuclei isolation

For the samples from therian species we developed a nuclei preparation method that includes fixation with dithio-bis(succinimidyl propionate) (DSP; or Lomant’s Reagent), a reversible cross-linker that stabilizes the isolated nuclei. The method was adapted from protocols used for the fixation of single cell suspensions^54^ and for the isolation of single nuclei from archived frozen brain samples^55^. Tissue pieces weighing ∼5 mg were homogenized in 100-150 µL/mg of prechilled lysis buffer (250 mM sucrose, 25 mM KCl, 5 mM MgCl_2_, 10 mM HEPES pH 8, 1% BSA, 0.1% IGEPAL, and freshly added 1 µM DTT, 0.4 U/µL RNase Inhibitor (New England BioLabs), 0.2 U/µL SUPERasIn (ThermoFischer Scientific)) and lysed for 5 min on ice. The lysate was centrifuged at 100 x g for 1 min at 4 °C. The supernatant was transferred to a new reaction tube and centrifuged at 500 x g for 5 min at 4 °C. The supernatant was removed and the pellet resuspended in 0.67 Vol (of volume lysis buffer used) of freshly made fixation solution (1 mg/mL DSP in PBS) and incubated for 30 min at room temperature. The fixation was quenched by addition of Tris-HCl to a final concentration of 20 mM. The fixed nuclei were pelleted at 500 x g for 5 min at 4 °C. The supernatant was removed and the pellet resuspended in 0.67 Vol of Wash Buffer (250 mM sucrose, 25 mM KCl, 5 mM MgCl_2_, 10 mM TRIS-HCl pH 8, 1% BSA, and freshly added 1 µM DTT, 0.4 U/µL RNase Inhibitor, 0.2 U/µL SUPERasIn). This was centrifuged at 500 x g for 5 min at 4 °C. The supernatant was removed and the pellet resuspended in 0.5 Vol of PBS. Then nuclei were strained using 40 µm Flowmi strainers (Sigma). For estimation of nuclei concentration, Hoechst DNA dye was added and the nuclei were counted using Countess™ II FL Automated Cell Counter (ThermoFischer Scientific).

For platypus and chicken, a similar preparation method was used, but the nuclei were not fixed. In brief, tissue pieces weighing ∼5 mg were homogenized in 100-150 µL/mg of prechilled lysis buffer (250 mM sucrose, 25 mM KCl, 5 mM MgCl_2_, 10 mM TRIS-HCl pH 8, 1% BSA, 0.1% IGEPAL, and freshly added 1 µM DTT, 0.4 U/µL RNase Inhibitor, 0.2 U/µL SUPERasin) and lysed for 5 min on ice. The lysate was centrifuged at 100 x g for 1 min at 4 °C. The supernatant was transferred to a new reaction tube and centrifuged at 500 x g for 5 min at 4 °C. The supernatant was removed and the pellet resuspended in 0.67 Vol of Wash Buffer. This was centrifuged at 500 x g for 5 min at 4 °C. The supernatant was removed and the pellet resuspended in 0.5 Vol of PBS. Then nuclei were strained using 40 µm Flowmi strainers (Sigma). For estimation of nuclei concentration, Hoechst DNA dye was added and the nuclei were counted using Countess™ II FL Automated Cell Counter (ThermoFischer Scientific).

### Preparation of sequencing libraries

For the construction of single-nucleus RNA-seq libraries, Chromium Single Cell 3’ Reagent Kits (10x Genomics; v2 chemistry for human, chimpanzee, bonobo, gorilla, gibbon, macaque, marmoset, mouse and opossum; v3 chemistry for platypus and chicken) were used according to the manufacturer’s instructions. 15,000 to 20,000 nuclei were loaded per lane in the Chromium microfluidic chips and cDNA was amplified in 12 PCR cycles. Sequencing was performed with NextSeq 550 (Illumina) according to the manufacturer’s instructions using the NextSeq 500/550 High Output Kit v2.5 (75 Cycles) with paired-end sequencing (read lengths of read1: 26 bp, read2: 57 bp; index1: 8bp, ∼170 to 380 million reads per library for v2 chemistry; read lengths of read1: 28 bp, read2: 56 bp; Index1: 8 bp, ∼247 to 306 million reads per library for v3 chemistry) (Supplementary Table 2).

For bulk RNA-seq data generation, RNA was extracted using the RNeasy Micro kit (QIAGEN). The tissues were homogenized in RLT buffer supplemented with 40 mM DTT. The RNA-seq libraries were constructed using the TruSeq Stranded mRNA LT Sample Prep Kit (Illumina) as described in ref.^4^. Libraries were sequenced on Illumina NextSeq550 using a single-end run (read1: 159 bp; index 1: 7 bp) with ∼24 to ∼60 million reads per library (Supplementary Table 2).

### Genome and transcript isoform annotation

Given that the quality of genome annotation differs substantially between the studied species and given the specific and widespread transcription of the genome in the testis^56^, we refined and extended previous annotations from Ensembl^28^ based on testis RNA-seq data. Specifically, akin to the procedure described in references^4,8^, we used previous extensive stranded poly(A)-selected RNA-seq datasets^4,57^ (100 nt, single-end) for human, macaque, mouse, opossum, platypus, and chicken, and generated and utilized stranded poly(A)-selected RNA-seq datasets (159 nt, single-end) for chimpanzee, gorilla, gibbon and marmoset. For each species, we downloaded the reference genome from Ensembl release 87 (ref. ^28^): hg38 (human), CHIMP2.1.4 (chimpanzee), rheMac8 (rhesus macaque), C_jacchus3.2.1 (marmoset), mm10 (mouse), monDom5 (opossum), and galGal5 (chicken); from Ensembl release 96 (ref. ^58^): gorGor4 (gorilla) and Nleu_3.0 (gibbon); and from Ensembl release 100 (ref. ^59^): mOrnAna1.p.v1 (platypus). Raw reads were first trimmed with cutadapt v1.8.3 (ref. ^60^) to remove adapter sequences and low-quality (Phred score < 20) nucleotides, then reads shorter than 50 nt were filtered out (parameters: -- adapter=AGATCGGAAGAGCACACGTCTGAACTCCAGTCAC --match-read-wildcards -- minimum-length=50 -q 20). Processed reads were then mapped to the reference transcriptome and genome using Tophat2 v2.1.1 (ref. ^61^) (parameters: --bowtie1 --read-mismatches 6 --read-gap-length 6 --read-edit-dist 6 --read-realign-edit-dist 0 --segment-length 50 --min-intron-length 50 --library-type fr-firststrand --max-insertion-length 6 --max-deletion-length 6). Next, we assembled models of transcripts expressed using StringTie v1.3.3 (ref. ^62^) (parameters: -f 0.1 -m 200 -a 10 -j 3 -c 0.1 -v -g 10 -M 0.5). Stringent requirements on the number of reads supporting a junction (-j 3), minimum gap between alignments to be considered as a new transcript (-g 10) and fraction covered by multi-hit reads (-M 0.5) were used to avoid merging independent transcripts and to reduce the noise caused by unspliced or incompletely spliced transcripts. We compared the assembled transcript models to the corresponding reference Ensembl annotations using the cuffcompare program v2.2.1 from the cufflinks package^63^. Finally, we combined the newly identified transcripts with the respective Ensembl gene annotation into a single gtf file. We extended the original Ensembl annotations by 21-61 Mbp with novel transcripts and by 23-49 Mbp with new splice isoforms (Supplementary Table 1).

### Raw reads processing

CellRanger v3.0.2 was used for platypus and chicken, and CellRanger v2.1.1 for the other species in line with the used Chromium chemistry. The CellRanger *mkref* function was used with default settings to build each species reference from genomic sequences and customized extended annotation files (Supplementary Table 1). Given that pre-mRNA transcripts are abundant in nuclei^64^, exons and introns features were concatenated as described in the CellRanger v2.1.1 documentation for considering intronic and exonic reads for gene expression quantification. The CellRanger *count* function was used with default settings to correct droplet barcodes for sequencing errors, align reads to the genome, and count the number of Unique Molecular Identifiers (UMI) for every gene/barcode combination.

### Identification of usable droplets

We used a customized approach for detection of usable droplets. This was done to account for the low RNA content of nuclei compared to the whole cells. Specifically, we used a knee point-based approach combined with the fraction of intronic reads as a marker of pre-mRNA transcripts (abundant in the nucleus), and *MALAT1* (nuclear-enriched long non-coding RNA) expression as a marker of nuclei (when present in the genome).

### Quality control of filtered cells

For each sample independently, high quality nuclei were selected removing outliers based on the number of UMIs and the percentage of mitochondrial RNA. We created a Seurat^65^ object using the Seurat R package v3.1.4 from the subset raw UMI count table generated by CellRanger corresponding to the usable droplets identified upstream, normalized the data using the *NormalizeData* function, identified the top 10,000 most variables genes using the *FindVariableFeatures* function, scaled the data using the *ScaleData* function, performed the PCA using the *RunPCA* function and calculated the Louvain clusters using the *FindNeighbors* (parameters: dims = 1:20) and *FindClusters* (dims = 1:20, resolution = 0.5) functions. The testis cell types are very diverse, and using global thresholds to eliminate outlier cells would remove entire cell types. Therefore, we filtered out outlier droplets for each cluster independently with values lower than the first quartile (Q1) - 1.5 × IQR (InterQuartile Range) and higher than the third quartile (Q3) + 1.5 × IQR for both the UMI content and the fraction of mitochondrial RNA. Then, we removed potential doublets using *doubletFinder_v3* function of DoubletFinder^66^ v2.0.1 (parameters: PCs = 1:20, pN = 0.25, nExp = 5% of the total number of cells, identifying pk using *paramSweep_v3, summarizeSweep* and *find*.*pK* functions).

### Orthologous gene sets

Gene expression comparisons across species were done based on 1:1 orthologous genes across the studied species, which were extracted from Ensembl^28^ using the biomaRt R package v2.40.5.

### Integration of datasets

From the previously filtered UMI count tables, we created Seurat objects for every sample independently, normalized the data and identified the top 10,000 most variable genes. Next, for each species independently, we applied the Seurat^65^ anchoring approach using *FindIntegrationAnchors* and *IntegrateData* functions with 20 principal components to integrate all datasets together into a single Seurat object correcting for the batch effect. For each integrated species-specific Seurat object, we normalized (*NormalizeData* function) and scaled the data (*ScaleData* function) and performed a PCA (*RunPCA* function). Louvain clusters were calculated using *FindNeighbors* and *FindClusters* functions (parameters: dims = 1:20, 1:20, 1:20, 1:20, 1:20, 1:20, 1:20, 1:17, 1:8, 1:10, 1:10 and resolution = 0.5, 0.5, 0.2, 0.5, 0.5, 0.5, 0.5, 0.5, 0.5, 0.3, 0.5, respectively, for human, chimp., bonobo, gorilla, gibbon, macaque, marmoset, mouse, opossum, platypus and chicken). The UMAP embedding coordinates were calculated using the *RunUMAP* function (parameters: dims = 1:20, 1:20, 1:20, 1:20, 1:20, 1:20, 1:20, 1:17, 1:10, 1:10, 1:10 and min_dist = 0.3, 0.3, 0.1, 0.1, 0.3, 0.3, 0.3, 0.1, 0.2, 0.3, 0.6 respectively for human, chimp., bonobo, gorilla, gibbon, macaque, marmoset, mouse, opossum, platypus and chicken).

All primate datasets were merged together using the LIGER^67^ (v0.5.0) integration tool. A LIGER object was created using the *createLiger* function based on primate 1:1 orthologues from Ensembl release 87, and normalized with *normalize, selectGenes* and *scaleNotCenter* functions with default settings. Then, the joint matrix was factorized using the *optimizeALS* function (k = 20) and the quantile normalization was performed with the *quantile_norm* and default settings. The Louvain clusters were calculated with the *louvainCluster* function and default settings as well as UMAP coordinates with the *runUMAP* function (n_neighbors = 100, min_dist = 0.2).

### Estimation of expression levels and normalization

The gene UMI counts per cell were normalized using the Seurat R package and its *NormalizeData* function. Therefore, the UMI counts of each gene in each cell are divided by the total UMI counts of each cell, multiplied by 10,000 and log-transformed.

### Cell type assignment

We identified the major cell type populations from the primate integrated, mouse, opossum, platypus and chicken objects independently using known marker genes^68-70^ mostly from human and mouse and their respective 1:1 orthologues in the other species. *CLU* marks Sertoli cells; *TAGLN* and *ACTA2* peritubular cells; *CD34* and *TM4SF1* endothelial cells; *APOE* and *CD74* macrophages; *STAR* and *CYP11A1* Leydig cells; *GFRA1, PIWIL4* (undifferentiated), *DMRT1*(differentiated), and *STRA8* spermatogonia; *SYCE1* (leptotene), *SYCP1* (zygotene), *PIWIL1* (pachytene), *SYCP2, TANK*, and *AURKA* spermatocytes; *LRRIQ1* (early), *ACRV1*, and *SPACA1* (late) round spermatids; *SPATA3, NRBP1, PRM1*, and *GABBR2* elongated spermatids. Cell type assignment was robustly reinforced by complementary analyses and metrics such as UMAP coordinates, pseudotime trajectories, transcriptional activities (UMI counts) and prior knowledge.

### Pseudotime

Pseudotime trajectories were calculated using the slingshot v1.2.0 R package^71^. We applied the *getLineages* function with the upstream calculated clusters and UMAP embedding coordinates of the germ cells to obtain connections between adjacent clusters using a minimum spanning tree (MST). We provided the starting and ending clusters based on the previous cell type assignment with known marker gene expression. Then we applied the *getCurves* function to the obtained lineages to construct smooth curves and order the cells along a pseudotime trajectory. Pseudotime values are highly variable depending on used tools, thus we ordered the cells one by one based on their pseudotime values and divided their rank by the total number of cells, to obtain evenly distributed values between 0 and 1. Finally, we validated the obtained pseudotime trajectories based on previous cell type assignments, expression patterns of marker genes and UMAP embedding coordinates.

### Marker gene identification

To precisely identify marker genes along spermatogenesis, we grouped the germ cells into 20 evenly distributed bins along the pseudotime trajectory for each species. Then, we applied the *FindAllMarkers* function (parameter: only.pos = T, min.pct = 0.25, logfc.threshold = 0.25, return.thresh = 0.05) of the Seurat v3.1.4 R package to the 22 groups (20 germline groups, the Sertoli and other somatic cell groups) (Extended Data Fig. 1; Supplementary Table 4).

### Global patterns of gene expression differences across mammals

Pseudo-bulk samples were generated using the *AverageExpression* function of the Seurat R package with various groups of cells depending on the pseudo-bulk samples produced in the study. For the analyses presented in Fig. 1b, we performed the principal component analysis (PCA) of normalized expression in amniote testicular cell types (pseudo-bulks) for each individual based on 4,498 1:1 amniote orthologues. PCA was performed using the *prcomp* function of the stats R package. For Fig. 1c, we constructed gene expression trees (as described in ref.^3^) using the neighbor-joining approach, based on pairwise expression distance matrices between corresponding pseudo-bulk samples for the different cell types across species. The distance between samples was computed as 1 – *ρ*, where *ρ* is Spearman’s correlation coefficient and was computed using the *cor* function of the stats R package. The neighbor-joining trees were constructed using the ape R package. The reliability of branching patterns was assessed with bootstrap analyses (the 4,498 1:1 amniote orthologues were randomly sampled with replacement 1,000 times). The bootstrap values are the proportions of replicate trees that share the branching pattern of the majority-rule consensus tree shown in the figures (Fig. 1c; Extended Data Fig. 2). The total tree length was calculated by removing the intra-species variability between individuals (Fig. 2a).

### Evolutionary forces

In Fig. 2c, we plotted the median pLI score across expressed genes (≥1 UMI) in each nucleus. We obtained the pLI scores from ref.^72^. For Fig. 2d, we used a set of neutrally ascertained knockouts consisting in 4,742 protein-coding genes, 1,139 of which are classified as lethal. For each cell, the denominator is the number of genes expressed that were tested for lethality and the numerator the genes among those that resulted in a lethal phenotype. Tested genes for viability and associated phenotype information were downloaded from the International Mouse Phenotyping Consortium (IMPC)^32^. For Fig. 2e (and Extended Data Fig. 3a), we used the average dN/dS values across 1:1 orthologues in primates. For each cell, the mean dN/dS value is plotted. Conserved 1:1 orthologues across 6 primates (human, chimpanzee, gorilla, gibbon, macaque and marmoset) as well as their coding and protein sequences were extracted from Ensembl^28^, providing a set of 11,791 protein-coding genes. For each species and orthologue the longest transcript was extracted. Orthologous protein sequences were aligned using clustalo v1.2.4; then pal2nal v14 was used (with protein sequences alignments and coding sequences as input) to produce codon-based alignments. The codeml software from the PAML package^73^ v4.9 was used to estimate dN/dS ratios. The M0 site model was applied to the orthologue alignments to estimate one average dN/dS ratio per orthologous gene set across species (parameter: NSites = 0, model = 0). In Fig. 2f (and Extended Data Fig. 3b), we plotted the percentage of positively selected genes expressed across nuclei. For each nucleus, the denominator is the number of expressed genes that were tested for signatures of positive selection and the numerator is the number of genes among those with evidence for positive selection. We used sets of genes previously identified as carrying evidence for coding-sequence adaptation in primates^11^ (331 positively selected genes out of 11,170 genes tested) and mammals^74^ (544 positively selected genes out of 16,419 genes tested).

In Fig. 2g (and Extended Data Fig. 3c), we plotted the average phylogenetic age of expressed genes across somatic and germ cells. The phylogenetic age of genes is an index that gives greater weight to young duplicates (as described in ref.^4,75^). The range of the score differs between species depending on the number of outgroup lineages available (more lineages allowed for more details in the phylogeny) and therefore this index cannot be compared across species, only within species (that is, across cells and cell types). The phylogenetic age of genes was obtained from GenTree (http://gentree.ioz.ac.cn/) with Ensembl release 69 (ref. ^75^). In Fig. 2h (and Extended Data Fig. 3d), we plotted the percentage of the cell transcripts originating from protein-coding genes and intergenic elements. Gene biotypes were obtained from Ensembl. Intergenic elements are all elements that are not protein-coding genes (lncRNA, pseudogenes, pseudogenes, and other sequences). In Fig. 2i (and Extended Data Fig. 3e), normalized log_2_-transformed median expression values across replicates at the transcriptome (e^*tr*^) and translatome (*e*^*tl*^) layers were used to calculate translation efficiency (TE = log_2_(e^*tr*^) – log_2_(e^*tl*^)) in testis (as described in ref.^8^) from RNA-seq and Ribo-seq, respectively. TE values were calculated at the whole testis level, thus only cell type-specific genes (for which 60% of their transcripts at the whole testis level are concentrated in a single cell type) were used. Higher values show a more efficient translation of transcripts, while lower values indicate relative translational repression. For Fig. 2j (and Extended Data Fig. 3f, g), we used time- and tissue-specificity indexes of expressed genes across somatic and germ cells in testis. As described in ref.^4^, tissue and time specificity indexes are calculated from RNA-seq data across organs and developmental stages. Both indexes range from 0 (broad expression) to 1 (restricted expression). The indexes were obtained from ref.^4^. For each nucleus, we plotted the median index across expressed genes. In Fig. 2k, we plotted the percentage of genes causing infertility when knocked-out (out of 3,252 knockouts, 173 of which caused infertility). Tested genes for infertility and associated phenotype information were downloaded from the IMPC database^32^. For each nucleus, the denominator corresponds to the number of genes expressed that were tested for infertility and the numerator to the genes among those that resulted in an infertility phenotype.

### Gene expression trajectories along spermatogenesis

We compared gene expression trajectories along spermatogenesis across primates using human, chimpanzee, bonobo, gorilla, gibbon, macaque and marmoset (based on Ensembl 87 orthologues), and across amniotes using human (as a representative of primates), mouse, opossum, platypus and chicken (based on Ensembl 100 orthologues). To compare robustly expressed genes, we used genes that are expressed in at least 5% of the cells in at least one cluster in all considered species. We used the mfuzz package^76^ (v.2.44.0), an unsupervised soft clustering method, to cluster gene expression trajectories along spermatogenesis (eight cell types in primates; four cell types in amniotes) across species using *Dmin* and *mestimate* functions to estimate the number of clusters and the fuzzification parameter (Extended Data Fig. 4). As described in ref.^4^, we inferred within a phylogenetic framework the probability that there were changes in trajectories along spermatogenesis, that is, that genes changed their cluster assignment in specific branches, using a 5% probability cutoff.

### Trajectory conservation score

We calculated a global trajectory conservation score across species for each 1:1 orthologous gene set. For a given orthologous gene set, this score corresponds to the log-transformed probability that all members fall into the same mfuzz trajectory cluster as:

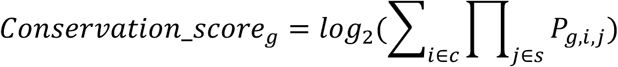

where *g* corresponds to a given orthologous gene set, *c* to all mfuzz trajectory clusters (1 to 9 for primates, 1 to 12 for amniotes), *s* to all species (human, chimp., bonobo, gorilla, gibbon, macaque, and marmoset for primates; human, mouse, opossum, platypus, and chicken for amniotes), and *P*_*g,i,j*_ to the probability that the gene g of the species j falls into the cluster i. A higher conservation score indicates a greater global trajectory conservation. As a proof of concept, we plotted the trajectory conservation score for conserved and changed trajectories which revealed a significant higher conservation score for conserved trajectories (Extended Data Fig. 7a).

### RNA *in situ* hybridization and expression quantification

Fresh testicular tissue was fixed in GR-fixative (7.4% formaldehyde, 4% acetic acid, 2% methanol, 0.57% Sodium Phosphate, Dibasic, and 0.11% Potassium Phosphate, Monobasic) overnight (for at least 16h) at 4 °C, dehydrated and embedded in paraffin. The *in situ* hybridization experiments were carried out on 4 µm sections mounted on SuperFrost Plus™ Slides (ThermoFisher Scientific) using the RNAscope 2.5 HD Detection Reagent RED kit according to the manufacturer’s recommendations (Advanced Cell Diagnostics). Briefly, testicular tissue sections were dewaxed in xylene and washed in 100% ethanol followed by treatment with hydrogen peroxide for 10 min. Target retrieval was performed for 15 or 30 min (see Supplementary Table 6 for specifications for each probe and species) using a steamer, followed by treatment with protease plus for 30 min at 40 °C. The slides were hybridized with the target probe (Supplementary Table 6) for 2 h at 40 °C followed by a series of signal amplifications (with amplification round 5 for 30 or 60 min). The sections were counterstained with Mayer’s hematoxylin and mounted with Vectamount® Permanent Mounting Medium (Vector Laboratories). The negative control probe *DapB* (a bacterial RNA) was run in parallel with the target probes and showed ≤5% positive cells in each section.

For *ADAMTS17* and *MYO3B*, positive (i.e., with red dots) round spermatids and spermatogonia were counted. For each section (human n=3, chimpanzee n=1, orangutan n=1), 10 tubules were counted using the NDP.viewPlus software (Hamamatsu Photonics). Two independent observers (S.B.W. and K.A.) counted positive and negative round spermatids and spermatogonia. No discrimination in intensity of the dots or the number of dots per cell was performed. Cell type identification was performed based on nucleus morphology and localization in the tubule. Only spermatogonia lining the edge of the tubules were counted (Extended Data Fig. 6a). The inter-observer variance was found to be 7% and 9% for round spermatids and spermatogonia, respectively. For *RUBCNL*, quantification of staining intensity was performed with R version 3.6.1 using countcolors^77^ (0.9.1) and colordistance^78^ (1.1.1) packages. For each section (human n=3, chimpanzee n=1, orangutan n=1), 10 tubules were divided into three parts: area dominated by spermatogonia, area dominated by spermatocytes (also containing Sertoli cells), and area dominated by spermatids (no distinction between the different types of spermatids) (Extended Data Fig. 6b). In each tubular area, the number of cells was counted manually using the NDP.view2Plus software (2.8.24). Then, the pixels occupied by red staining were quantified and the expression level for each cell type was calculated by dividing the stained pixels by the number of cells. For each picture, the stained pixels for each cell type were normalized by the total amount of stained pixels.

### Cell type and testis-specific genes per chromosome

Testis-specific genes were obtained from previously generated RNA-seq data^4^ of adult organs (RPKM ≥1 in testis and RPKM < 1 in brain, cerebellum, heart, kidney, and liver). Among these, cell-type specific genes were studied for each chromosome. Somatic cell type specific genes were identified using the *FindAllMarkers* function (parameter: only.pos = TRUE, min.pct = 0.05, logfc.threshold = 0.25, return.thresh = 0.05). Germ cell type specific genes were identified using mfuzz trajectory clusters previously calculated. We first calculated the percentage of genes located on a given chromosome among all genes in the genome (x-axis of plots in Extended Data Fig. 8a, c; red horizontal line for X-linked genes in Fig. 4b). We then contrasted this with the percentage of testis-specific genes with predominant expression in a given cell type (y-axis of plots in Extended Data Fig. 8a,c; y-axis of plots in Fig. 4b for X-linked genes). Finally, the percentage of testis-specific genes per cell type and chromosome was statistically compared to the percentage of genes per chromosome in the genome using exact binomial tests.

### Classification of X and Y-bearing spermatids

The Y chromosome carries a low number of genes and is missing in some genome assemblies. Thus, we focused on the fraction of X transcripts in spermatids to classify them as X or Y-bearing cells. For this, we fitted a Gaussian Mixture Model to the data with 2 components (bimodal distribution) independently for each replicate, using the function *normalmixEM* of the mixtools (v.1.2.0) R package. The two obtained normal distributions were utilized to classify X (higher levels of X transcripts) and Y-bearing (lower levels of X transcripts) spermatids using 95% confidence intervals. Outlier and overlapping cells were not assigned to either category. Finally, we checked that the fraction of Y transcripts was significantly higher in Y-bearing spermatids (Fig. 4c). Bifurcating UMAPs (Fig. 4c) were obtained using X- and Y-linked genes in addition to previously identified highly variable genes to perform the PCA associated with the UMAP coordinate calculation.

### General statistics and plots

Unless otherwise stated, all statistical analyses and plots were done in R v3.6.2 (ref.^79^). Plots were created using ggplot2 v3.2.1, tidyverse v1.3.0, cowplot v1.0.0 and pheatmap v1.0.12.

## Data availability

Raw and processed bulk and single-nucleus RNA-seq data have been deposited in ArrayExpress with the accession codes E-MTAB-11063 (human snRNA-seq), E-MTAB-11064 (chimpanzee snRNA-seq), E-MTAB-11067 (bonobo snRNA-seq), E-MTAB-11065 (gorilla snRNA-seq), E-MTAB-11066 (gibbon snRNA-seq), E-MTAB-11068 (macaque snRNA-seq), E-MTAB-11069 (marmoset snRNA-seq), E-MTAB-11071 (mouse snRNA-seq), E-MTAB-11072 (opossum snRNA-seq), E-MTAB-11070 (platypus snRNA-seq), E-MTAB-11073 (chicken snRNA-seq) and E-MTAB-11074 (chimpanzee, gorilla, gibbon and marmoset bulk RNA-seq) (https://www.ebi.ac.uk/arrayexpress/). All other data are available as supplementary information or available upon request. The testis gene expression at the single-nucleus level across the eleven studied species can be visualized using the shiny app we developed: https://apps.kaessmannlab.org/SpermEvol/.

## Code availability

Custom code will be made available upon request.

## Supplementary Tables

The file contains Supplementary Tables 1-8 and a guide for the supplementary tables.

## Extended Figures

**Extended Data Fig. 1.**
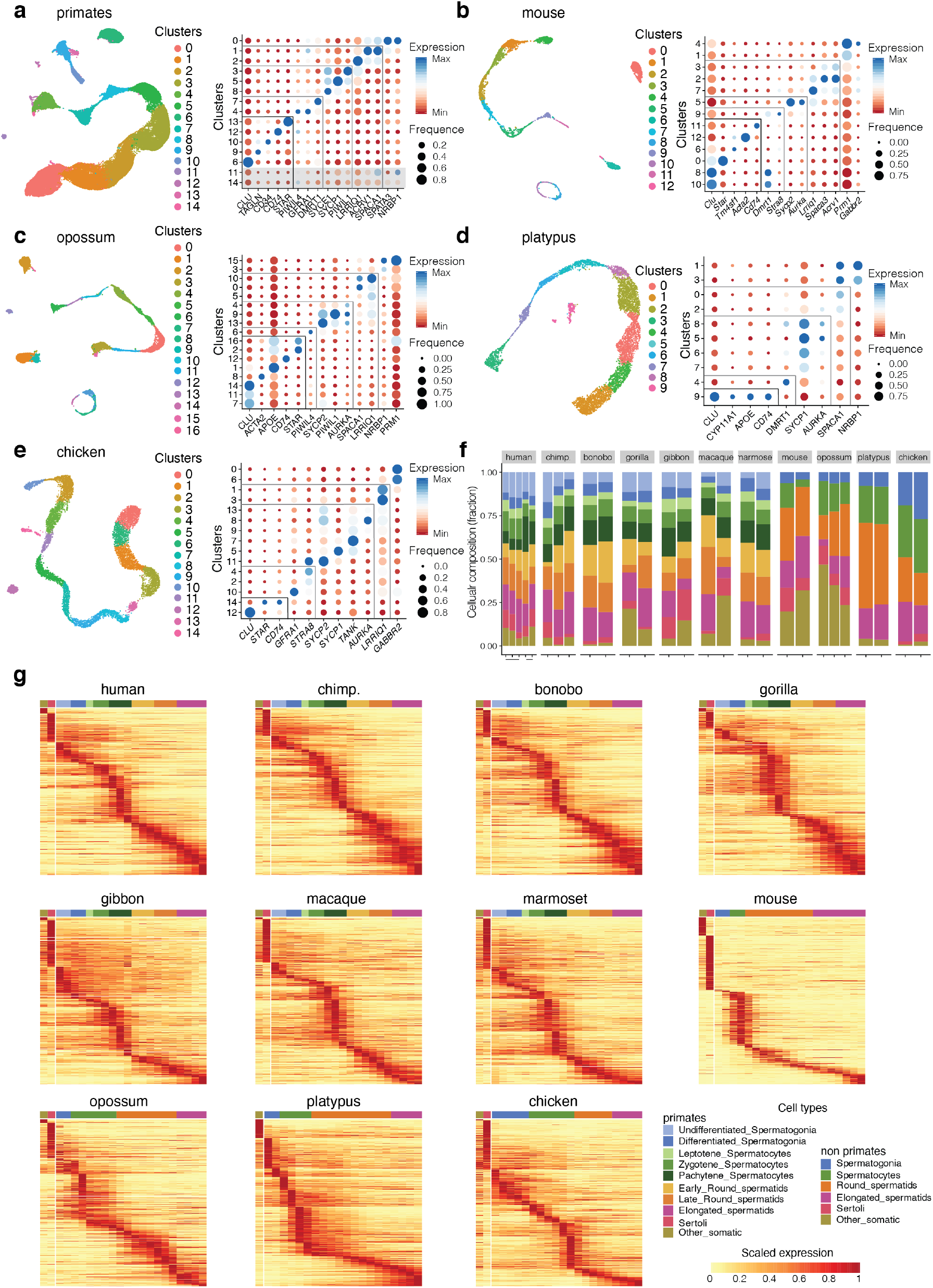
Cellular composition and marker gene assessment. **a-e**, Clustering and UMAP representations (left), marker gene-based cell type assignment (right), for **a**, primates, **b**, mouse, **c**, opossum, **d**, platypus and **e**, chicken. *CLU* marks Sertoli cells; *TAGLN* and *ACTA2* peritubular cells; *CD34* and *TM4SF1* endothelial cells; *APOE* and *CD74* macrophages; *STAR* and *CYP11A1* Leydig cells; *GFRA1, PIWIL4* (undifferentiated), *DMRT1*(differentiated), and *STRA8* spermatogonia; *SYCE1* (leptotene), *SYCP1* (zygotene), *PIWIL1* (pachytene), *SYCP2, TANK* and *AURKA* spermatocytes; *LRRIQ1* (early), *ACRV1* (late) and *SPACA1* (late) round spermatids; *SPATA3, NRBP1, PRM1* and *GABBR2* elongated spermatids. **f**, Cell-type composition. **g**, Scaled expression of specific genes (y-axis) for all species and cell types (x-axis).

**Extended Data Fig. 2.**
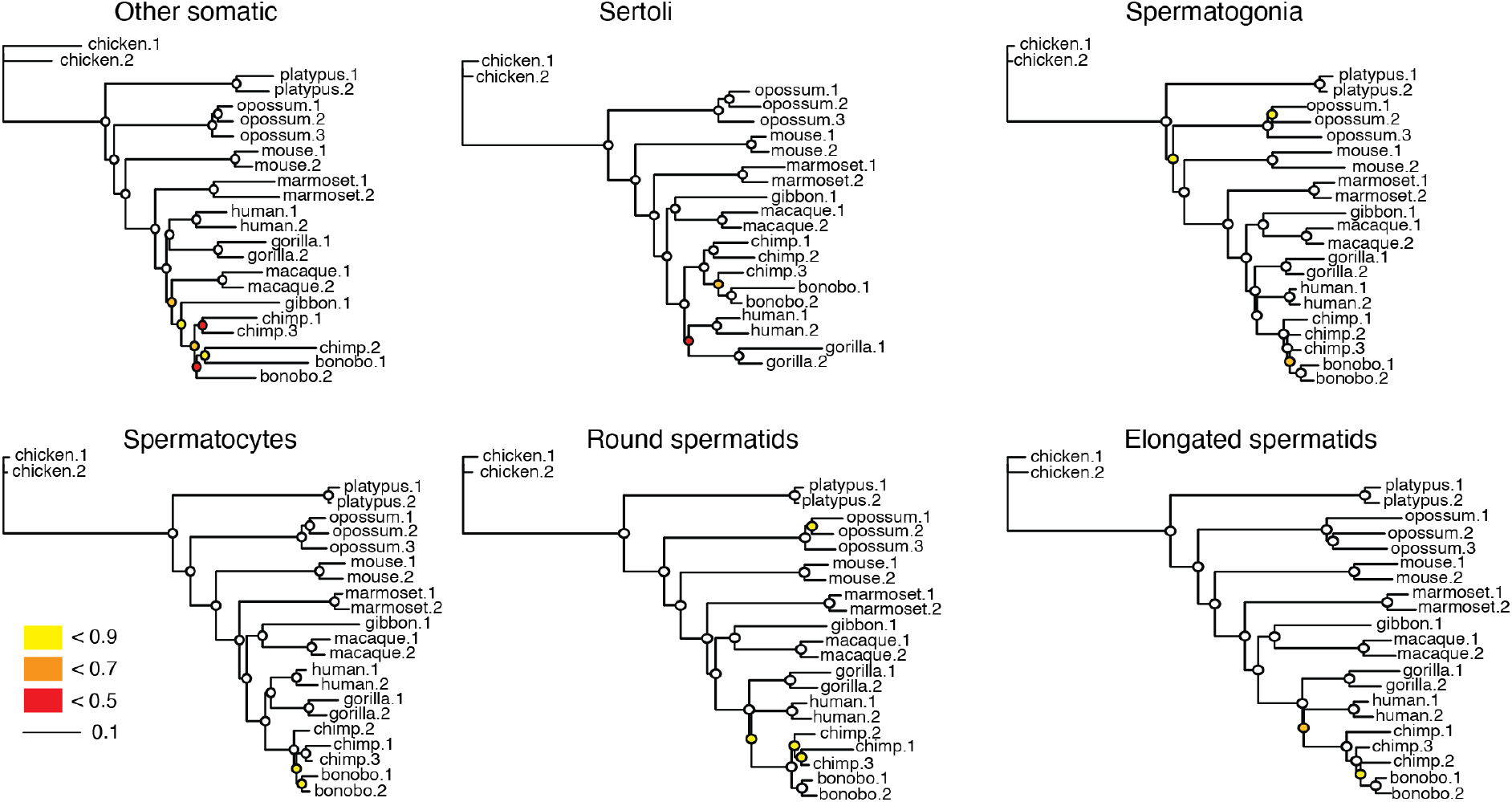
Mammalian testicular gene expression phylogenies. Neighbor-joining trees based on pairwise expression level distances (1-rho, Spearman’s correlation coefficient) for the main testicular cell types, respectively. Trees are drawn to the same scale (indicated by the scale bar). Bootstrap values (i.e., proportions of replicate trees that have the branching pattern as in the majority-rule consensus tree shown) are indicated by circles at the corresponding nodes: ≥ 0.9 (white fill).

**Extended Data Fig. 3.**
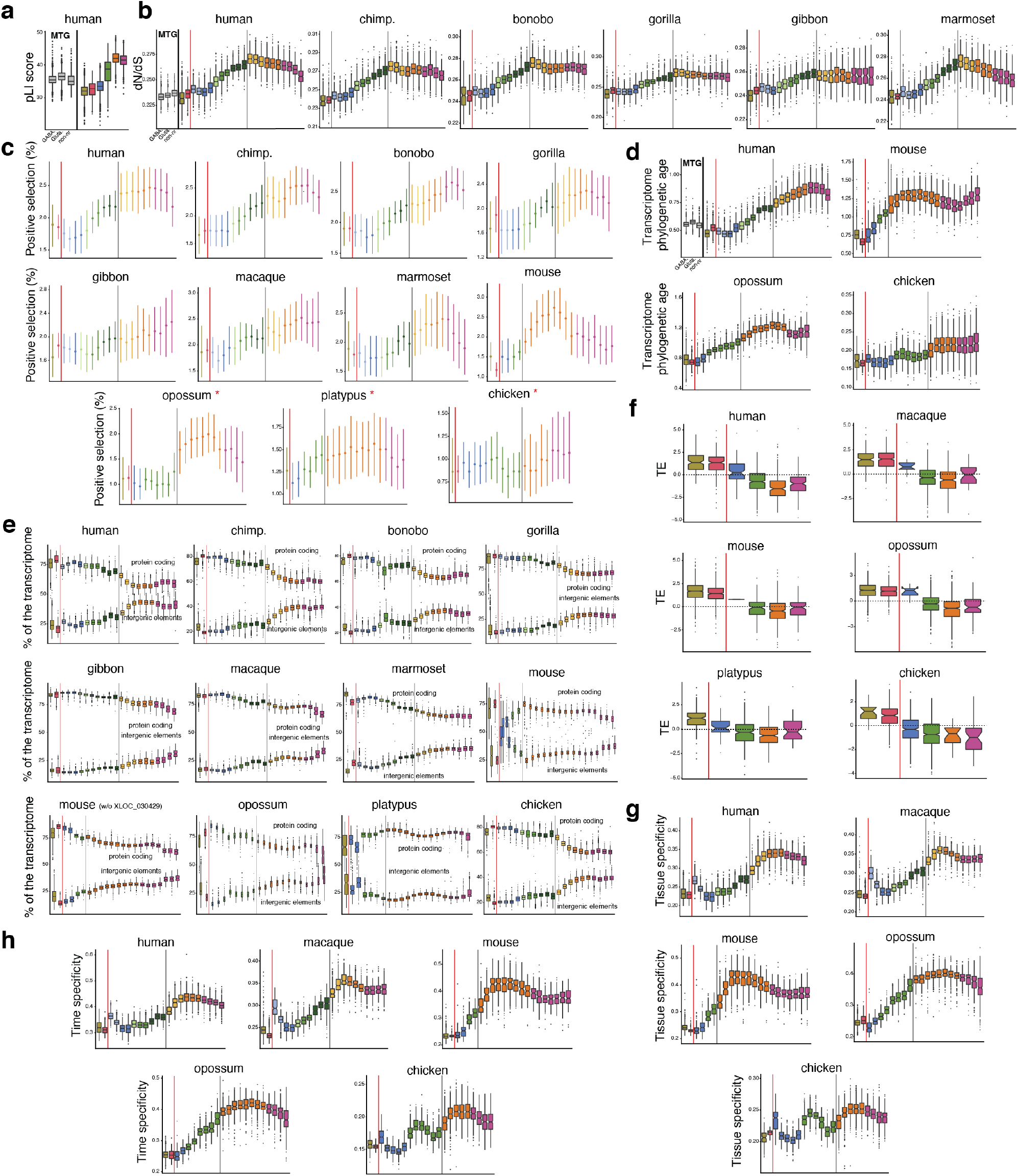
Evolutionary forces. **a**, Tolerance to functional variants. **b**, Average normalized ratio of nonsynonymous over synonymous nucleotide substitutions (dN/dS) of expressed genes across primates. **c**, Percentage of positively selected genes along spermatogenesis. Opossum, platypus, and chicken were not used in the original studies^11,74^ and so the 1:1 orthologues in these species were used. **d**, Phylogenetic age of cellular transcriptomes. Higher values reflect increased lineage-specific gene contributions (that is, younger transcriptomes). **e**, Percentages of UMIs that map to protein coding genes (top) or to intergenic elements (bottom). XLOC_030429 drives a different mouse pattern. **f**, Translational efficiency (TE; normalized log2-transformed values) of expressed genes. **g**, Tissue specificity. **h**, Time specificity (development). In **a, b**, and **d**, human middle temporal gyrus (MTG) snRNA-seq data^80^ was used for non-gonadal (brain) comparisons.

**Extended Data Fig. 4.**
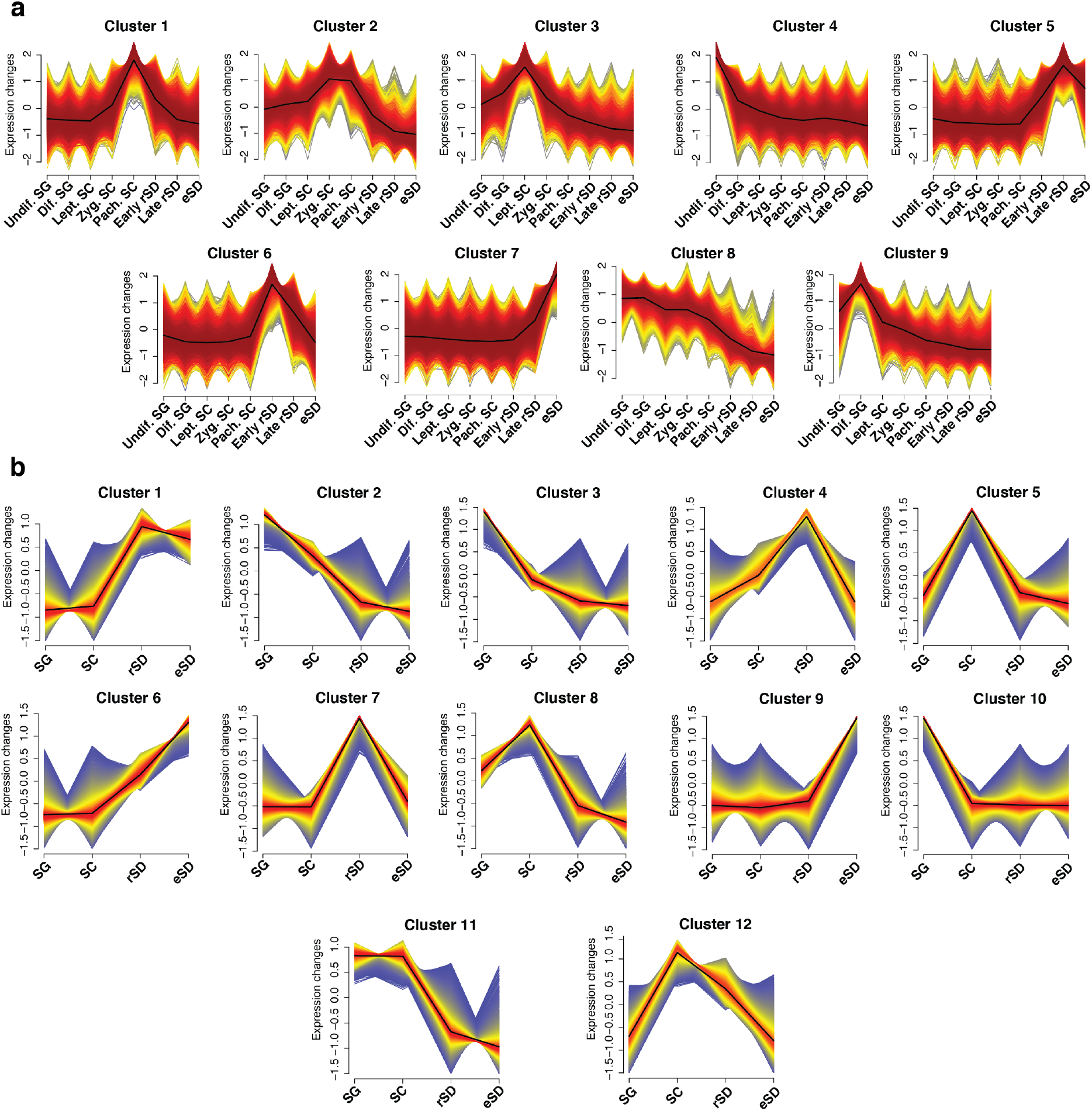
Gene expression trajectory clusters. **a**, Gene expression trajectory clusters for primates (human, chimp., bonobo, gorilla, gibbon, macaque, marmoset). **b**, Gene expression trajectory clusters for amniotes (human, mouse, opossum, platypus, chicken).

**Extended Data Fig. 5.**
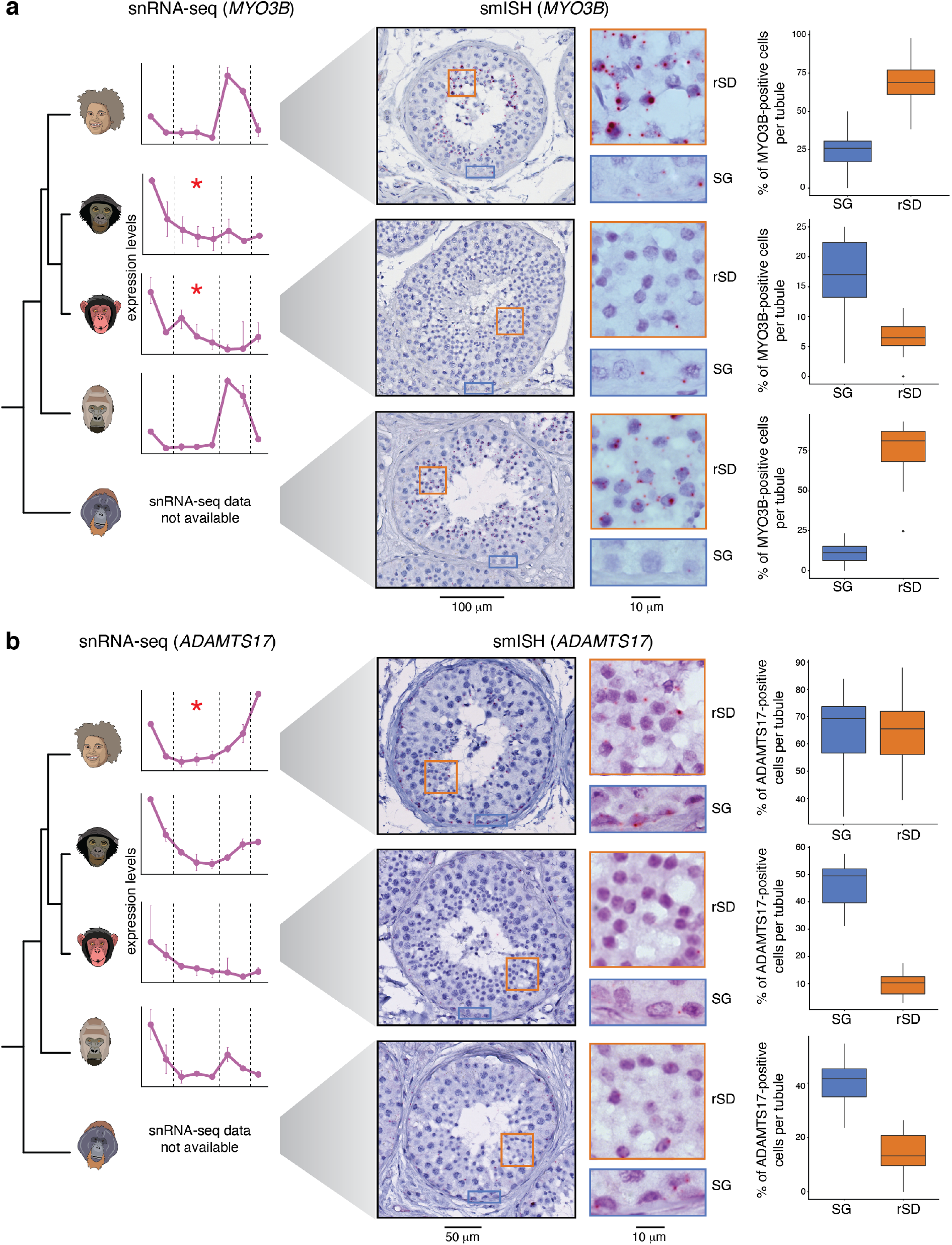
Examples of gene expression trajectory changes along primate spermatogenesis validated by smISH. Left: gene expression trajectory from snRNA-seq along spermatogenesis for human, bonobo, chimp., and gorilla (from top to bottom). Center: images of seminiferous tubule cross sections from human, chimp., and orangutan (from top to bottom) stained with smISH for *MYO3B* (**a**) and *ADAMTS17* (**b**) using RNAScope. Red asterisks indicate on which branch a trajectory change was called. Examples of spermatogonia (SG) and round spermatids (rSD) closeups are provided for visualization purposes. Right: box plots represent the percentage of positive cells in SG and rSD from 10 tubules per section (n=3 for human, n=1 for chimp. and orangutan) with whiskers at 1.5 times the interquartile range.

**Extended Data Fig. 6.**
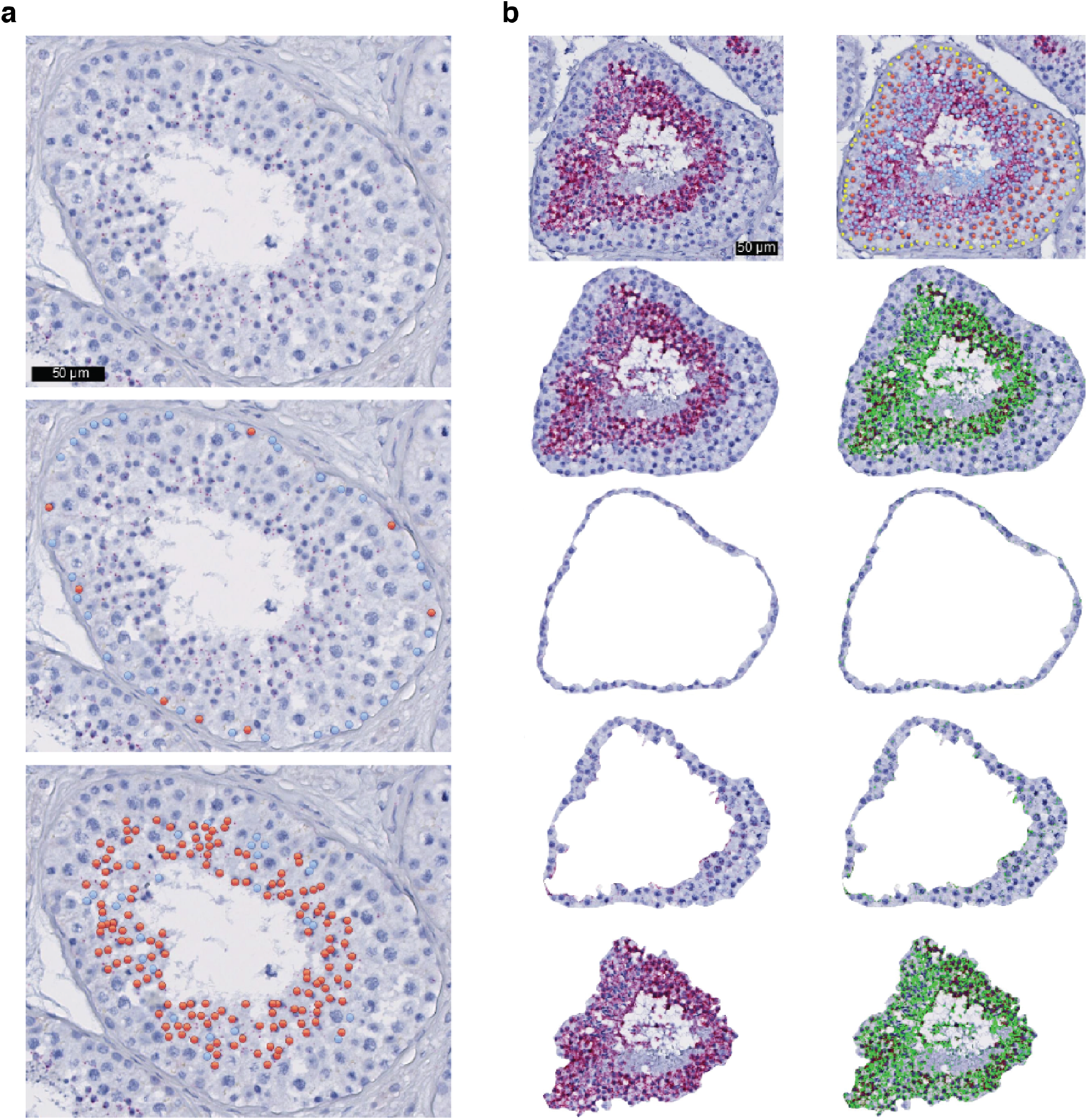
Quantification of gene expression from smISH experiments. **a**, Example of *MYO3B* staining and expression quantification in an orangutan seminiferous tubule. Dots show *MYO3B* positive (red) and negative (blue) spermatogonia (middle) and round spermatids (bottom). **b**, Example of *RUBCNL* staining and expression quantification in a chimpanzee seminiferous tubule. Single-molecule RNA staining is shown in red and computationally detected transcripts are colored in green. The first row shows *RUBCNL* staining (left) and the three main spermatogenic cell types (spermatogonia, in yellow; spermatocytes, in red; and spermatids, in blue). The second, third, fourth and fifth rows show *RUBCNL* staining (left) and detected transcripts (right) at the whole tubule, spermatogonia, spermatocytes and spermatids levels, respectively.

**Extended Data Fig. 7.**
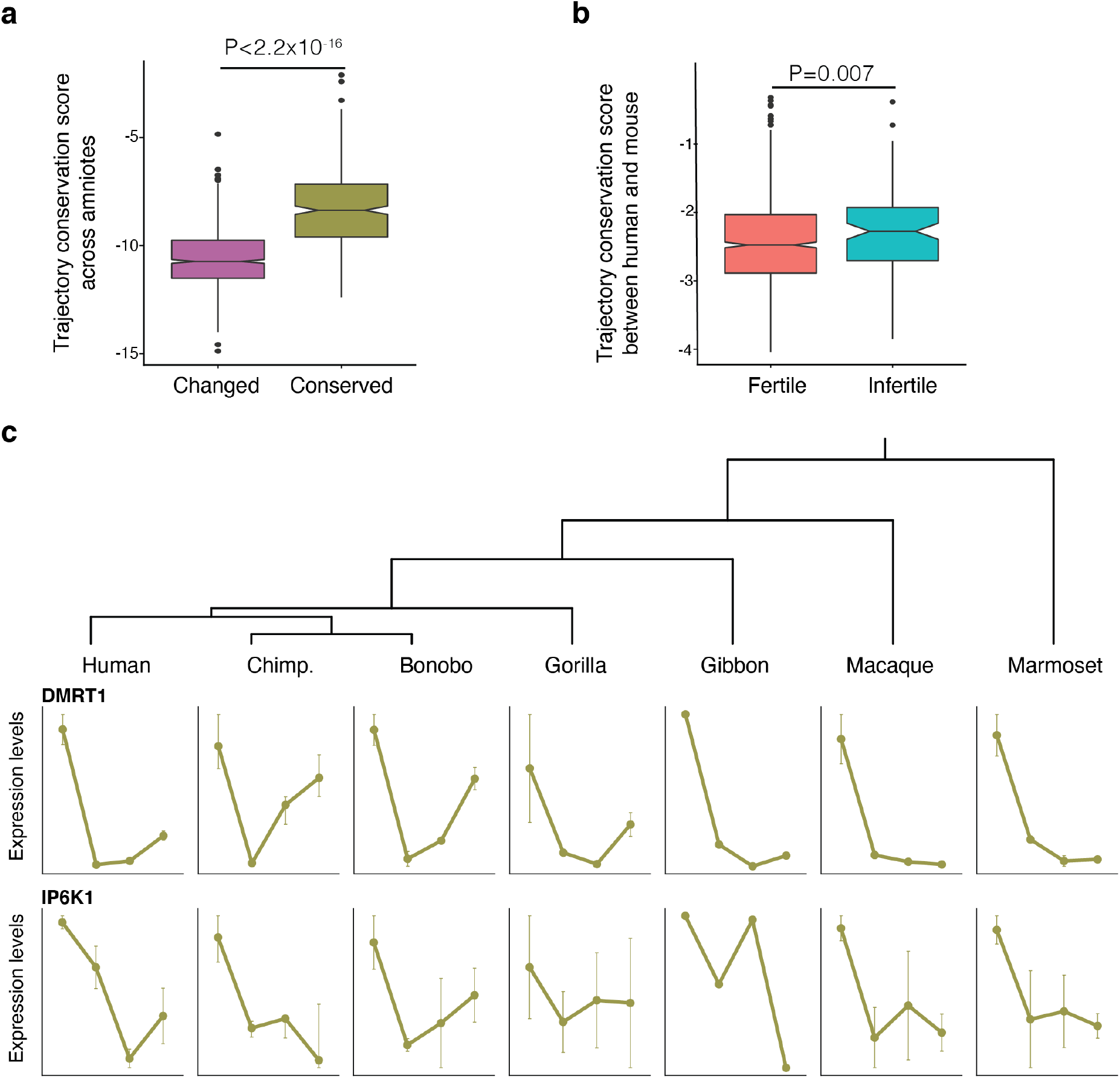
Conservation of gene expression trajectories along spermatogenesis. **a**, Global trajectory conservation score across amniotes for changed and conserved trajectories (*P* < 2.2×10^−16^, two-sided Wilcoxon rank-sum test). **b**, Trajectory conservation score between human and mouse of genes that lead to either a fertile or an infertile phenotype when knocked out in mouse (*P* = 0.007, two-sided Wilcoxon rank-sum test). **c**, Gene expression trajectories (*DMRT1*, top; *IP6K1*, bottom) along primate spermatogenesis (spermatogenesis, from left to right). Dots depict the median gene expression across replicates for each cell type and the whiskers show minimum and maximum gene expression values across replicates.

**Extended Data Fig. 8.**
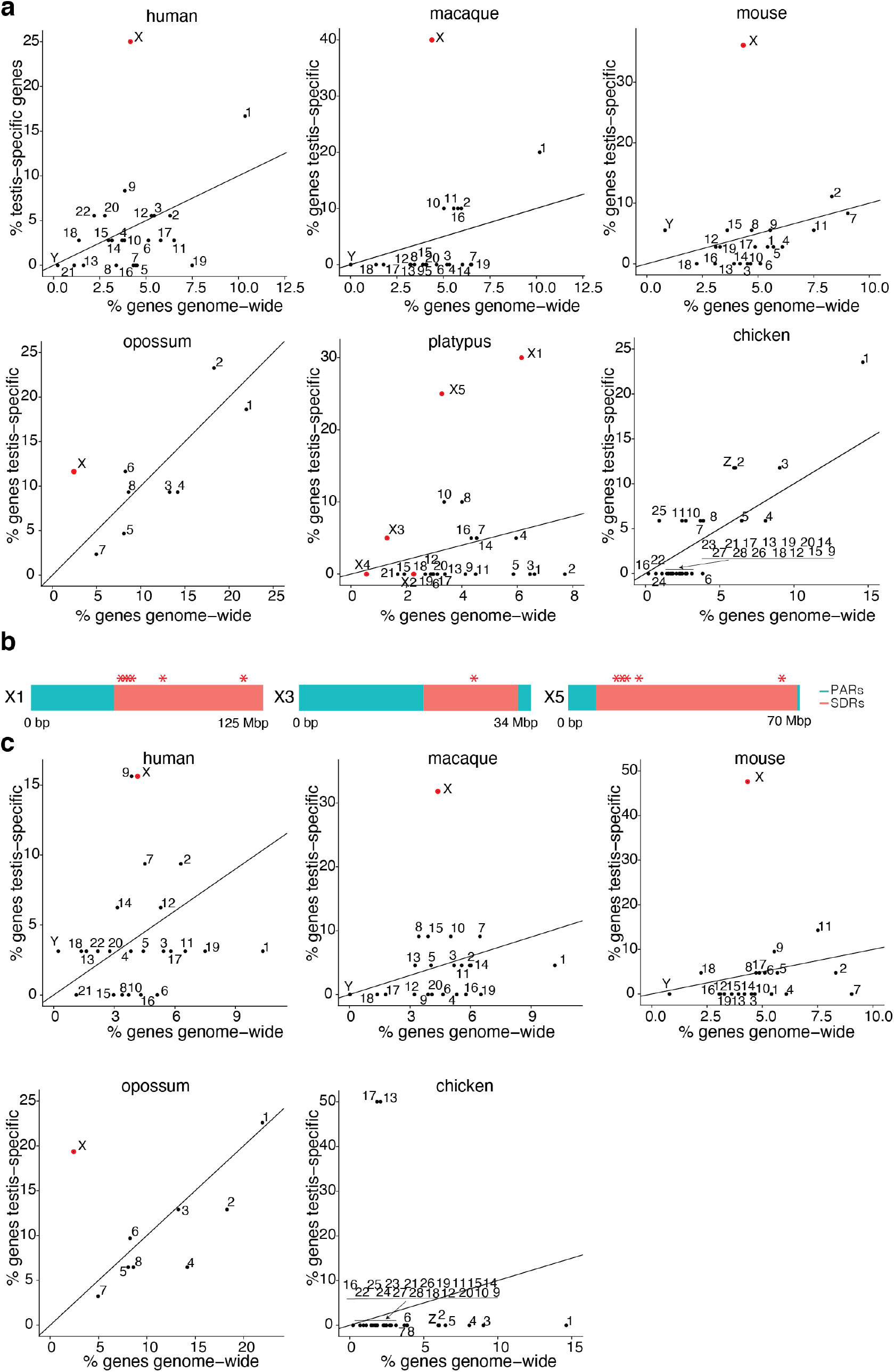
Per chromosome testis-specific genes enriched in spermatogonia and Sertoli cells. **a, c**, Per chromosome percentage of expressed testis-specific genes (y-axis) with predominant expression in differentiated spermatogonia (human and macaque) or spermatogonia (mouse, opossum and platypus) (**a**) and Sertoli cells (**c**) versus the percentage of all genes in the genome (x-axis). X chromosomes are colored in red. The diagonal shows the numbers expected if testis-specific genes were randomly distributed across the genome. **b**, Location of platypus testis-specific genes enriched in spermatogonia on chromosomes X1, X3 and X5.

**Extended Data Fig. 9.**
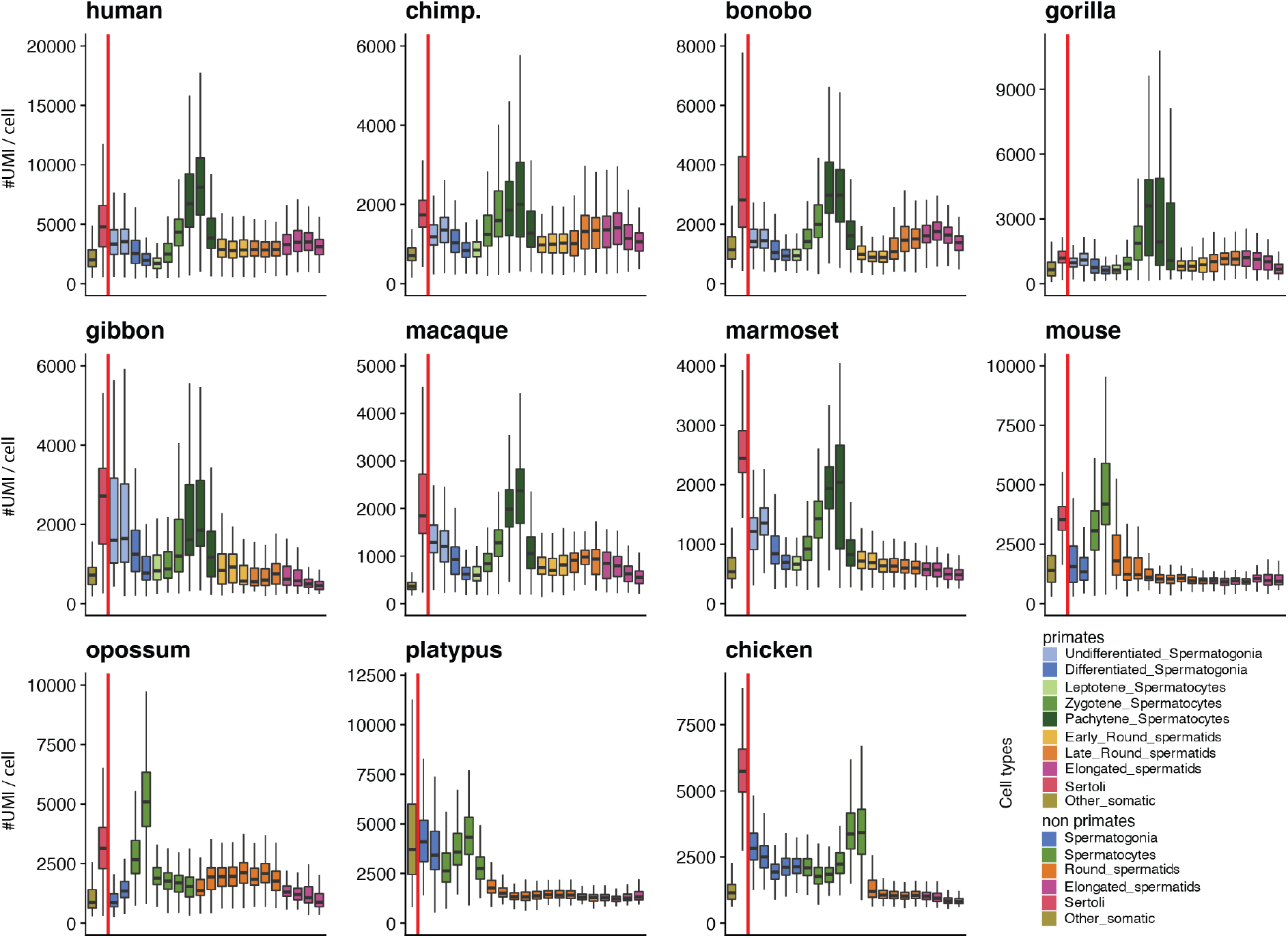
Transcription levels in testicular cell types. Number of UMIs per cell in somatic cells (left of the vertical red bar) and along spermatogenesis (right of the vertical red bar, from left to right) across species.

**Extended Data Fig. 10.**
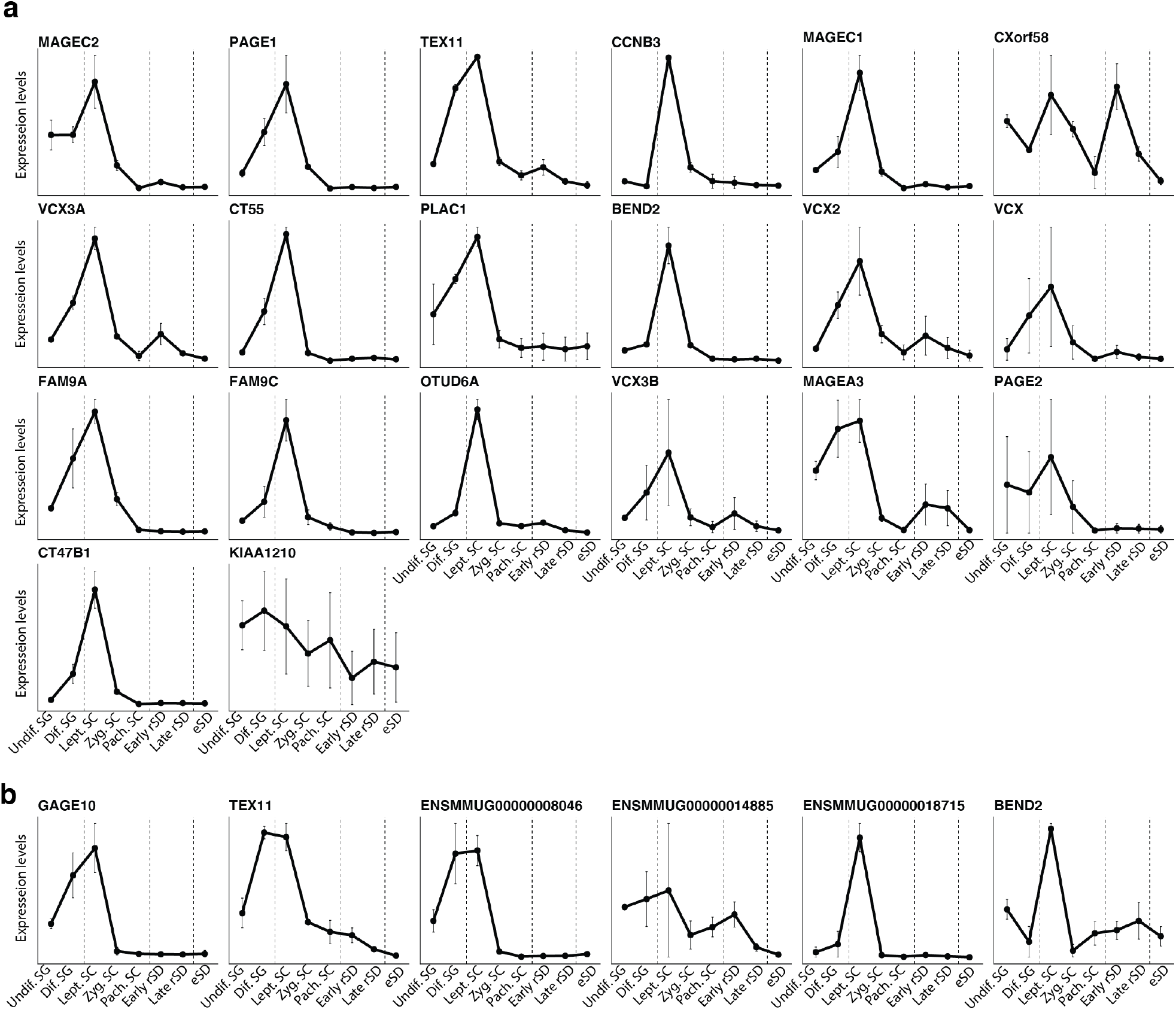
Trajectories. Trajectories of testis-specific X-linked genes with predominant expression in leptotene spermatocytes in human (**a**) and macaque (**b**) along spermatogenesis.

**Extended Data Fig. 11.**
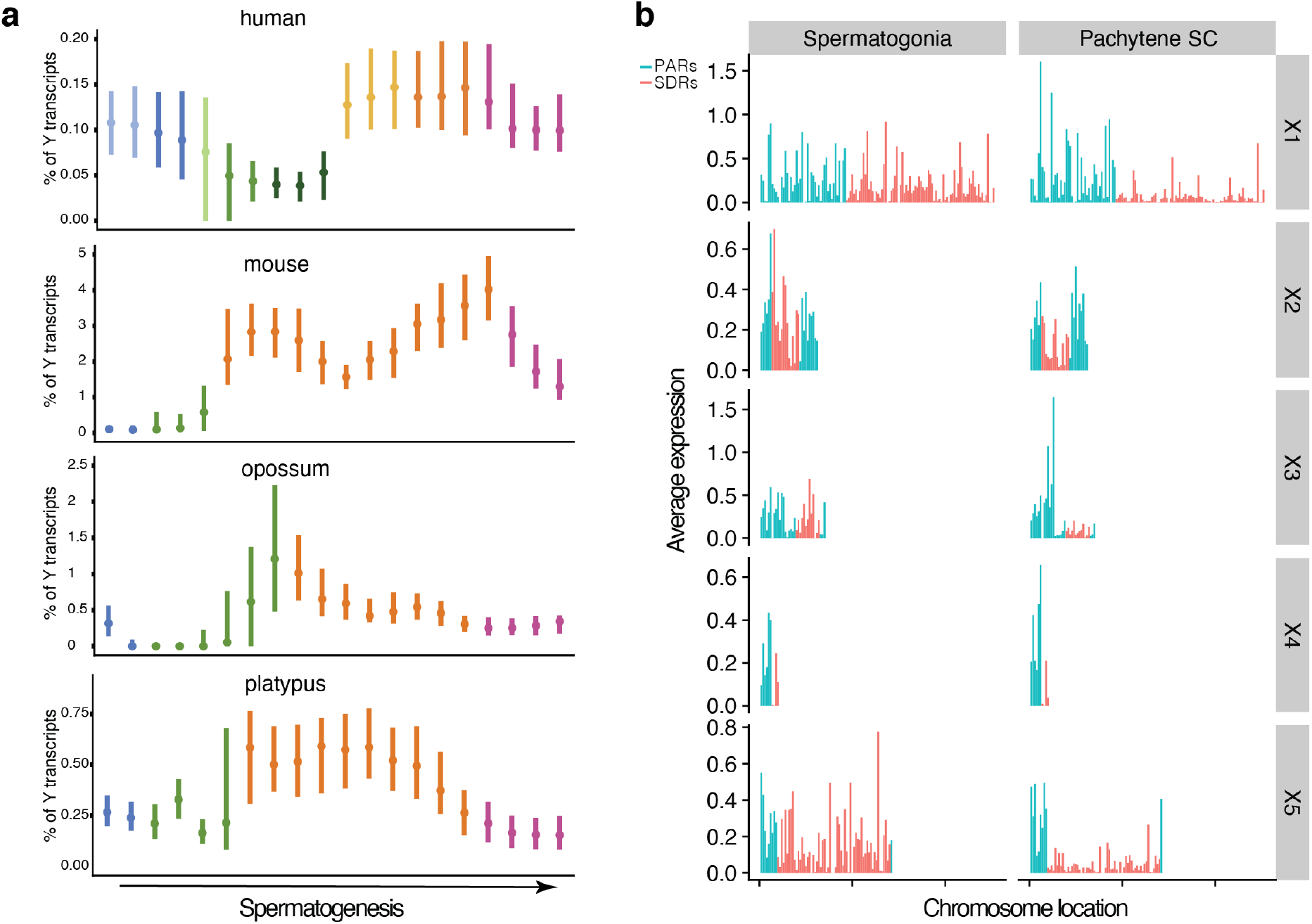
Mammalian sex chromosome evolution. **a**,Percentage of Y transcripts along spermatogenesis. For spermatids, only Y-bearing cells are considered. Germ cells are binned into 20 equally sized groups across spermatogenesis (from left to right). Reported values correspond to median values across genes expressed in each cell of the different testicular cell types. Whiskers depict the median ± 25th and 75th percentiles. **b**, Average expression along platypus X chromosomes in spermatogonia and pachytene spermatocytes. Genes are grouped into 1,000,000 bp bins along the chromosomes.

